# Macrophage mediated recognition and clearance of *Borrelia burgdorferi*nelicits MyD88-dependent and -independent phagosomal signals that contribute to phagocytosis and inflammation

**DOI:** 10.1101/593566

**Authors:** Sarah J. Benjamin, Kelly L. Hawley, Paola Vera-Licona, Carson J. La Vake, Jorge L. Cervantes, Rachel Burns, Oscar Luo, Yijun Ruan, Melissa J. Caimano, Justin D. Radolf, Juan C. Salazar

## Abstract

Lyme disease is a tick-borne illness caused by the spirochete *Borrelia burgdorferi* (Bb). It is believed that the robust inflammatory response induced by the host’s innate immune system is responsible for the clinical manifestations associated with Bb infection. The macrophage plays a central role in the immune response to many bacterial infections and is thought to play a central role in activation of the innate immune response to Bb. Previous studies have shown that following phagocytosis of spirochetes by macrophages, phagosome maturation results in degradation of *Bb* and liberation of bacterial lipoproteins and nucleic acids, which are recognized by TLR2 and TLR8, respectively, and elicit MyD88-mediated phagosome signaling cascades. Bone marrow-derived macrophages (BMDMs) from *MyD88*^−/−^ mice show significantly reduced spirochete uptake and inflammatory cytokine production when incubated with *Bb ex vivo*. Paradoxically, additional studies revealed that Bb-infected *MyD88*^−/−^ mice exhibit inflammation in joint and heart tissues. To determine the contribution of MyD88 to macrophage-mediated spirochete clearance, we compared wildtype (WT) and *MyD88*^−/−^ mice using a murine model of Lyme disease. *MyD88*^−/−^ mice showed increased Bb burdens in hearts 28 days post infection, while H&E staining and immunohistochemistry showed significantly increased inflammation and greater macrophage infiltrate in the hearts of *MyD88*^−/−^ mice. This suggests that Bb triggers MyD88-independent inflammatory pathways in macrophages to facilitate cell recruitment to tissues. Upon stimulation with Bb *ex vivo*, WT and *MyD88*^−/−^ BMDMs exhibit significant differences in bacteria uptake, suggesting that MyD88 signaling mediates cytoskeleton remodeling and the formation of membrane protrusions to enhance bacteria phagocytosis. A comprehensive transcriptome comparison in Bb-infected WT and *MyD88*^−/−^ BMDMs identified a large cohort of MyD88-dependent genes that are differentially expressed in response to Bb, including genes involved in actin and cytoskeleton organization (*Daam1, Fmnl1*). We also identified a cohort of differentially-expressed MyD88-independent chemokines (*Cxcl2, Ccl9*) known to recruit macrophages. We identified master regulators and generated networks which model potential signaling pathways that mediate both phagocytosis and the inflammatory response. These data provide strong evidence that MyD88-dependent and -independent phagosomal signaling cascades in macrophages play significant roles in the ability of these cells to phagocytose Bb and mediate infection.

**AUTHOR SUMMARY:** Macrophages play prominent roles in bacteria recognition and clearance, including *Borrelia burgdorferi* (Bb), the Lyme disease spirochete. To elucidate mechanisms by which MyD88/TLR signaling enhances clearance of *Bb* by macrophages, we studied Bb-infected wildtype (WT) and MyD88−/− mice and Bb-stimulated bone marrow-derived macrophages (BMDMs). Bb-infected MyD88−/− mice show increased bacterial burdens, macrophage infiltration and altered gene expression in inflamed heart tissue. MyD88−/− BMDMs exhibit impaired uptake of spirochetes but comparable maturation of phagosomes following internalization of spirochetes. RNA-sequencing of infected WT and MyD88−/− BMDMs identified a large cohort of differentially expressed MyD88-dependent genes involved in re-organization of actin and cytoskeleton during phagocytosis along with several MyD88-independent chemokines involved in inflammatory cell recruitment. We computationally generated networks which identified several MyD88-independent master regulators (*Cxcl2* and *Vcam1*) and MyD88-dependent intermediate proteins (*Rhoq* and *Cyfip1*) that are known to mediate inflammation and phagocytosis respectively. These results provide mechanistic insights into MyD88-mediated phagosomal signaling enhancing Bb uptake and clearance.

## INTRODUCTION

Lyme disease (LD) is a highly prevalent tick-borne illness caused by the spirochetal bacterium *Borrelia burgdorferi* (Bb) (1-3). The disease is characterized by a wide array of clinical manifestations which vary in duration and severity between patients. Early clinical manifestations of LD include the characteristic “bullseye” rash known as *erythema migrans* and flu-like symptoms, while late manifestations include arthritis, carditis and neurological compromise (4, 5). The invading spirochete induces both innate and adaptive immune responses, and it is believed that the innate immune response to Bb contributes to the development of the clinical findings characteristic of LD (6). The macrophage is a principal cellular element of the innate immune response to the bacterium at sites of infection in both humans and mice (7-9). Macrophages also play a prominent role in the pathogenesis of murine Lyme carditis, and their recruitment to heart tissue is important in spirochetal clearance (10). Macrophages have the phagocytic and signaling machinery necessary to bind, engulf, and degrade Bb. Binding of Bb to macrophages is mediated by surface integrins, such as Complement Receptor 3 (CR3) (11, 12) and *α*_3_ (13). Once attached, phagocytosis of Bb is complex and can occur by either a sinking or coiling mechanism (14, 15). Both cases require rearrangements of the actin cytoskeleton to internalize Bb into the endosome, where degradation takes place (16).

Bb is an extracellular pathogen that needs to be taken up and degraded for significant recognition by the host immune system (17). We have defined this process as “phagosomal signaling” (14). Spirochete degradation exposes borrelial <u>p</u>athogen-<u>a</u>ssociated <u>m</u>olecular <u>p</u>atterns (PAMPs), such as lipoproteins and nucleic acids, to endosomal toll-like receptors (TLRs) for recognition, resulting in signaling cascades which induce pro-inflammatory cytokine production (17-19). Bb does not contain LPS and, therefore, does not engage TLR4. The cell envelope of Bb contains abundant triacylated lipoproteins (20), which are known to be recognized by TLR1/2 heterodimers (21-26). However, the three fatty acid chains in the N-terminus of Bb lipoproteins, which serve as the TLR2/1 PAMP, are tethered in the outer membrane (27). We have shown that this results in minimal recognition of lipoproteins in intact spirochetes at the cell surface (17, 18, 21, 28). Instead, principal recognition of Bb TLR2 ligands occurs within macrophage endosomal structures after the spirochete is phagocytosed and degraded (17, 28). Bacterial degradation results in exposure of both lipoprotein ligands and nucleic acids, which are recognized by endosomal TLR2 and TLRs 7, 8 and 9 respectively (18, 19, 29). Signaling cascades initiated by engagement of these TLRs utilize the adaptor protein MyD88(14, 30), indicating that this adaptor protein is a crucial element in mediating the inflammatory response to Bb.

A role for MyD88 has been implicated in each of the four general steps associated with phagocytic clearance of bacterial pathogens: uptake, maturation, degradation and cytokine production. Murine macrophages lacking MyD88 show markedly diminished uptake of several bacterial species, including Bb (28, 31-35). Prior studies have shown that Bb-induced MyD88 signaling results in increased PI3K activation, while inhibition of PI3K results in decreased Bb uptake by macrophages (36). In addition, formin proteins (FMNL1, mDia1, and Daam1) have also been shown to play a critical role in mediating phagocytosis of Bb (15, 37). Whether MyD88 increases activation of these formins, and the role of PI3K signaling in this process, has not been established. Degradation of bacteria is also impaired in the absence of MyD88 due to inefficient acidification of phagosomes (38). In the context of Bb infection, lysosome maturation markers are recruited to Bb-containing phagosomes in macrophages lacking MyD88 (28). However, the degree of phagosome maturation and acidification required to expose Bb ligands from the bacteria cell envelope for recognition has not been studied. Murine macrophages lacking MyD88 also show markedly diminished production of NFκB-triggered pro-inflammatory cytokines, such as TNFα and IL-6, when stimulated with different bacterial species, including Bb (28, 31). Nevertheless, the key host components involved downstream of these MyD88-mediated phagosome signals and their effects have not been well studied in the context of Bb infection.

Here we show using a murine model of LD that MyD88 plays a very important role in carditis severity and enhances bacterial clearance *in vivo*. We also show that there are tissue-specific inflammatory transcriptional responses that correlate with increased macrophage infiltration in the absence of MyD88. Using an *ex vivo* murine macrophage system, we show that MyD88 signaling enhances, but is not required, for bacterial uptake or phagosomal maturation. Through RNA-sequencing analysis, we provide evidence that MyD88 signaling drives transcription of multiple genes involved in phagocytosis and identify potential intermediate proteins that facilitate the association between MyD88 and bacterial uptake. We also demonstrate that uptake of Bb by macrophages induces robust MyD88-independent inflammatory responses via phagocytosis-dependent production of chemokine proteins that can mediate cell recruitment to tissues. Our findings highlight the importance of MyD88 in efficient uptake of the Lyme disease spirochete by macrophages, and provide potential mechanistic insight into how MyD88 mediates this process.

## RESULTS

### MyD88 signaling significantly reduces bacterial load in tissues of Bb-infected mice

As previously stated, the macrophage is an essential cellular element of the human inflammatory response to the LD spirochete (7). Macrophages have also been shown as part of the inflammatory cell infiltrate in heart and joint tissue of mice experimentally infected with Bb (10, 39), and the importance of MyD88 in Bb clearance from mouse tissues has been previously reported (40-42). However, the effect of MyD88 signaling on macrophage-specific inflammation and spirochetal clearance has not been well characterized. To do so, we used a murine model of LD that enables investigation of the cell populations present in Bb-infected tissues, its transcriptional responses and bacterial clearance. For these experiments, we needle-inoculated WT and MyD88−/− mice on a C57BL/6 background with 1×10^5^ Bb and quantitated bacterial burdens in heart, tibiotarsal joints, patellofemoral joints, bladder, ear and skin tissues at 14-, 28- and 56-days post-infection (DPI). This expands on work done in prior studies by investigating burdens in multiple tissues at various time points. By 14 days, all experimentally infected mice had seroconverted (data not shown). In heart tissue, MyD88−/− mice showed significantly increased bacterial burdens compared to WT mice at 14 and 28 DPI (**Figure S1A**); this difference was not significant by 56 DPI. WT heart tissue showed decreased Bb burdens at each time point when compared to the other tissues analyzed (**Figure S1A**). In joint, bladder, ear and skin tissues, burdens in WT and MyD88−/− mice were comparable at 14 DPI, but significantly higher in MyD88−/− mice at 28 and 56 DPI (**Figure S1B-F**). Interestingly, we saw a significant decrease in bacterial burdens in WT mice in all tissues except heart between 14 and 56 DPI (**Figure S1B-F**). Taken together, our results substantiate prior studies that MyD88 signaling is essential for Bb control; our results also show that, in heart tissue, MyD88-independent mechanisms can initiate effective, though delayed, bacterial clearance which could partially be macrophage mediated.

### Murine macrophages are a major component of heart inflammatory infiltrate independently of MyD88

To determine whether MyD88 has an effect on macrophage-mediated inflammation in Bb-infected murine hearts, we first examined the cellular infiltrates at 14 and 28 DPI by hematoxylin and eosin (H&E) staining. The histopathology of infected hearts consisted of lymphocytes, plasma cells, macrophages, reactive mesothelial cells, and fibroblasts. In more intensely inflamed heart tissue, neutrophils were also a significant component of the cellular infiltrate (**Figure 1A**). Inflammation was primarily evident in connective tissue at the heart base, especially around the root of the aorta, and in the atrial epicardium with some involvement of the atrial myocardium, valve leaflets and ventricular epicardium (10, 43, 44). Similar numbers of WT (2/5) and MyD88−/− (2/5) infected mouse hearts had evidence of low-grade inflammation (scores of 0-1) by 14 DPI (**Figure 1A and 1B**). Infected WT hearts showed stromal cell thickening at the base (**Figure 1A, top panels**), whereas a neutrophil infiltrate was clearly evident in the heart of one MyD88−/− infected mouse 14 DPI (**Figure 1A, inset**). Importantly, inflammation scores between the two genotypes were not significantly different at 14 DPI (**Figure 1B**). By contrast, at 28 DPI, the majority of MyD88−/− mice (4/5) showed significantly increased cellular infiltrates (**Figure 1C, bottom panels**) and higher inflammation scores than WT mice (**Figure 1D**). Hearts from MyD88−/− mice also revealed the presence of neutrophils (**Figure 1C, bottom panels and inset**), whereas WT mice did not. (**Figure 1C, top panels**). Because macrophages are not easily distinguishable by H&E staining, we performed immunohistochemistry (IHC) analysis for Iba-1, a calcium channel marker known to be expressed in macrophages (45). At 14 DPI, small numbers of macrophage clusters were observed in the heart base in WT mice (**Figure 1E, top left**), while no increase in macrophages was observed in MyD88−/− hearts 14 DPI (**Figure 1E, top right**). By contrast, 28 DPI MyD88−/− mouse hearts revealed a very dense macrophage infiltrate at the base (**Figure 1E, bottom right**), while minimal macrophage infiltrate was observed in WT hearts (**Figure 1E, bottom left**). To correlate cell infiltrate with an inflammatory response, we performed RT-PCR of the *Tnfa* gene from total RNA isolated from heart tissues from Bb-infected mice. As shown in **Figure 1F**, there was a significant decrease in *Tnfa* transcription between WT and MyD88−/− mice at 14 DPI, with WT hearts showing marked upregulation compared to uninfected WT controls. However, by 28 DPI *Tnfa* transcript levels were reduced in WT mice and comparable to those of MyD88−/− mice (**Figure 1G**). Taken together, our findings suggest that although infiltration into heart tissue is delayed, murine macrophages are still significant components of murine Lyme carditis in the absence of MyD88.

**Figure 1:**
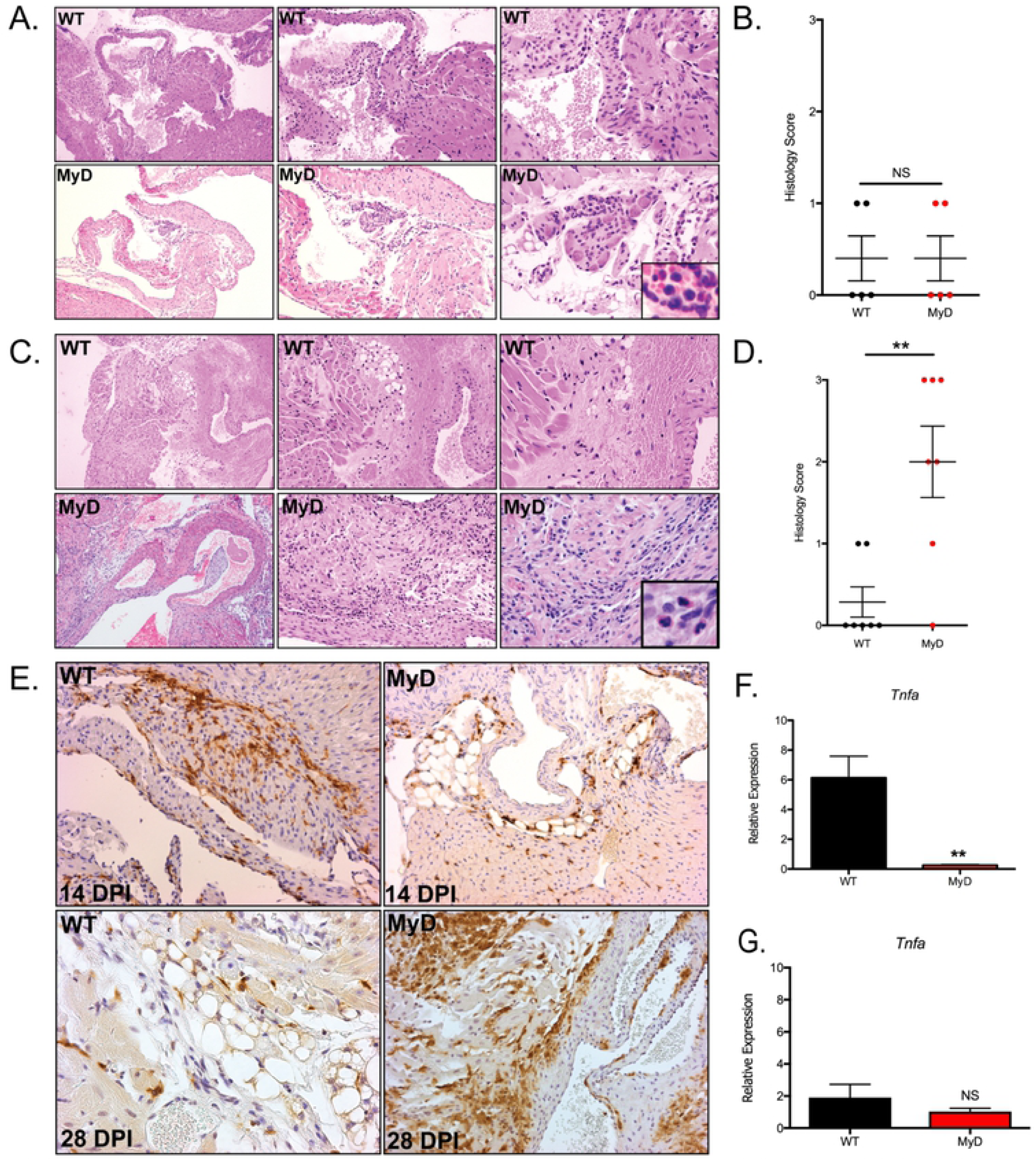
MyD88−/− mice show increased inflammation severity and macrophage infiltrate 28 days post Bb-infection. (A and C) Representative images of H&E staining of heart tissue sections from WT and MyD88−/− mice syringe-inoculated with 10^5^ Bb at 14 DPI (A) and 28 DPI (C). Images are of increasing magnification from left to right (10x, 20x, 40x, inset is 60x). Sections are 5μm. (B and D) Compilation of inflammation scores for WT (black dots) and MyD88−/− (red dots) mice at 14 DPI (B) and 28 DPI (D) N=5 mice per group. (E) Iba-1 immunohistochemistry of heart tissue sections from Bb-infected WT and MyD88−/− mice, magnification = 20x. Sections are 5μm. (F and G) Compilation of qRT-PCR *Tnfa* gene amplifications from heart tissue RNA. Total RNA was isolated from Bb-infected WT and MyD88−/− mice 14 (F) and 28 (G) days post infection. Gene amplification values were normalized to *Gapdh*. *p-value<0.05, **p-value<0.01, NS=not significant

### The absence of MyD88 affects expression of genes associated with inflammation but does not increase inflammation severity or infiltration of macrophages in joints

We performed similar analyses of patellofemoral joint tissue to evaluate macrophage responses in the absence of MyD88. Inflamed joints had small numbers of neutrophils, lymphocytes, plasma cells and macrophages (**Figure S2A and S2C**). Cellular infiltrates were most evident in the joint capsule near the junction with the periosteum. Prominent mixed infiltrates were also present in the periarticular loose connective tissue, especially in fat and along the periosteum, extending distal from the joint. A low-grade immune cell infiltrate was observed in most WT (3/5) and all MyD88−/− (5/5) mouse joints 14 DPI (**Figure S2A**). Similar to heart tissue, no differences in inflammation score between WT and MyD88−/− joints were observed 14 DPI (**Figure S2B**). The numbers of WT (3/5) and MyD88−/− (5/5) mice presenting with inflammation 14 DPI also were comparable (**Figure S2B**). In contrast to hearts, inflammation severity in joints was minimal to mild at 28 DPI in both genotypes, with no appreciable differences in the cell types present (**Figure S2C, bottom panels**). Inflammation scores also were comparable between WT and MyD88−/− mice; no significant difference in the number of mice showing inflammation was noted (WT = 2/5, MyD88−/− = 3/5) (**Figure S2D**). Using Iba-1 IHC, we visualized a few macrophages in the periarticular space in both WT and MyD88−/− mice 14 DPI (**Figure S2E, top panels**) and 28 DPI (**Figure S2E, bottom panels**). However, no difference in the number of macrophages was apparent between the two genotypes at the time points studied. There was also no difference in *Tnfa* transcript production at 14 DPI (**Figure S2F**) or 28 DPI (**Figure S2G**). This indicates that Bb induces tissue-specific inflammatory responses, even in the absence of MyD88 signaling.

### MyD88-deficient macrophages show comparable binding but reduced uptake of Bb

It has been well established that MyD88 enhances phagocytosis of multiple bacterial species by macrophages (28, 32, 34, 35, 46). To better understand the contribution of MyD88 to spirochete binding, uptake and degradation by macrophages, we transitioned into an *ex vivo* macrophage model using WT and MyD88−/− BMDMs co-incubated with Bb at MOIs of either 10:1 or 100:1 for 1, 4 or 6 hours. To quantify binding percentages, we imaged a minimum of 100 cells by confocal microscopy and from the images determined the number of cells with spirochetes either attached to the surface or internalized (**Figure 2A, yellow and white arrows respectively**). We used the same confocal images and total cell numbers to quantify uptake percentages based on the number of cells with internalized spirochetes. The percentages of cells with spirochetes either bound or internalized were comparable between WT and MyD88−/− BMDMs at all three time points irrespective of MOI (**Figure 2B and 2D**). While macrophages of both genotypes were able to phagocytose Bb, MyD88−/− BMDMs showed significantly reduced spirochete uptake compared to WT BMDMs (**Figure 2C**). Increasing the MOI to 100:1 significantly enhanced uptake in both cell genotypes, but MyD88−/− BMDMs never reached the phagocytic potential of their WT counterparts (**Figure 2E**). These results further support the necessity of MyD88 signaling for efficient phagocytosis of Bb, irrespective of contact time with the spirochete.

**Figure 2:**
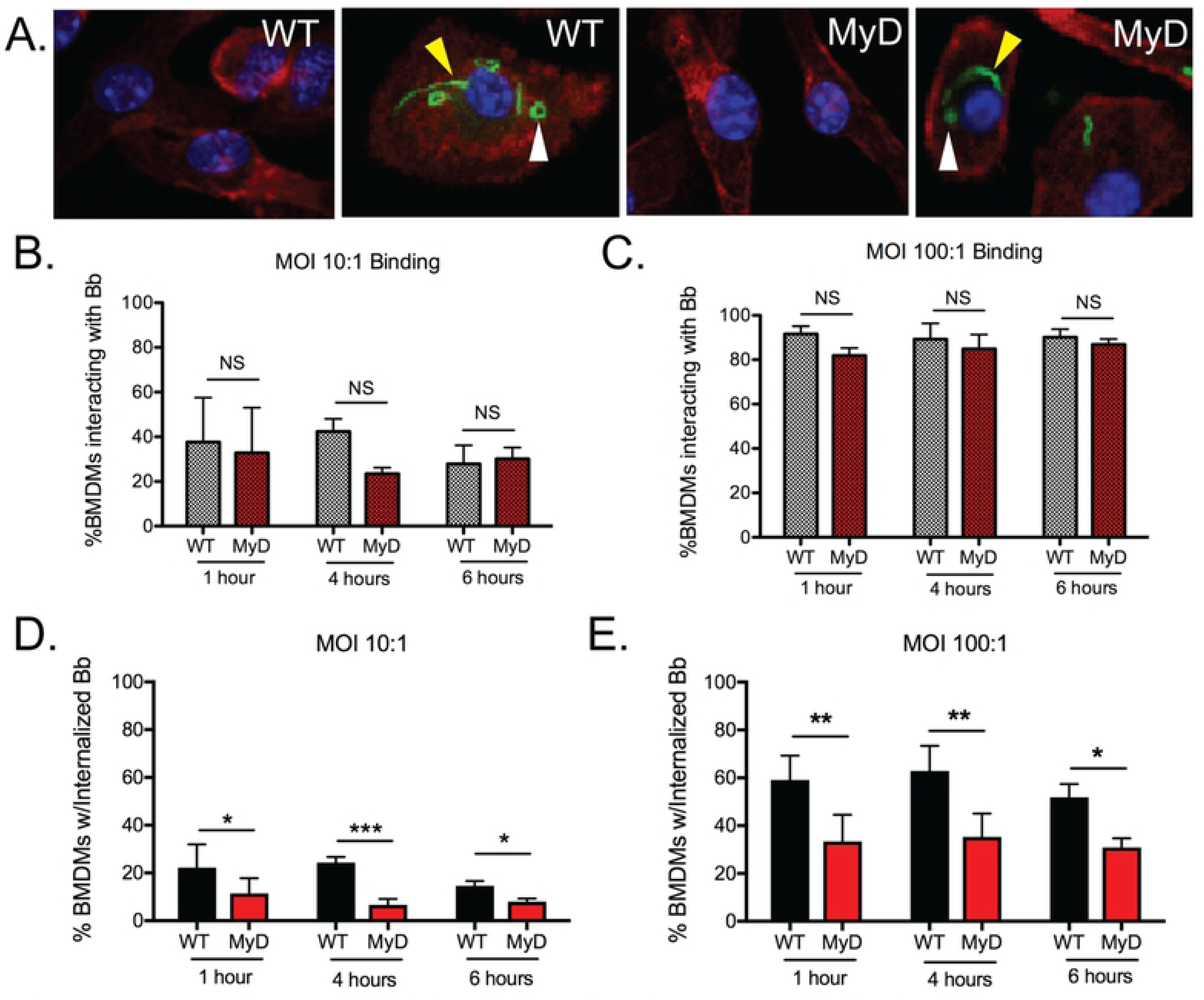
Quantitation of Bb binding and uptake by WT and MyD88−/− BMDMs. (A) Confocal images of WT and MyD88−/− BMDMs after 6 hours of stimulation with Bb at MOI 10:1, highlighting bound (yellow arrows) and internalized (white arrows) spirochetes. Green is Bb, red is actin and blue is cell nucleus. (B-C) Quantitation of bound spirochetes to WT (black bars) or MyD88−/− (red bars) BMDMs after 1, 4 or 6 hours of stimulation at a MOI of 10:1 (B) or 100:1 (C). (D-E) Quantitation of internalized spirochetes to WT (black bars) or MyD88−/− (red bars) BMDMs after 1, 4 or 6 hours of stimulation at MOI 10:1 (D) or 100:1 (E). n=3-5 mouse BMDM experiments per genotype *p-value<0.05, **p-value<0.01, ***p-value<0.001, NS=not significant

### TLR2, TLR7 and MyD88 are recruited to Bb-containing phagosomes in macrophages

Once spirochetes are phagocytosed by macrophages, recruitment of TLR and MyD88 proteins to the phagosome is essential to trigger MyD88-dependent signaling cascades (47-50). Importantly, we have demonstrated that in human monocytes TLR2 and TLR8 co-localize to endosomes containing Bb (19). In addition, other groups have shown a prominent role for TLR7 in the Bb inflammatory response (51). Murine TLR8, unlike murine TLR7 and human TLR8, does not seem to utilize ssRNA as its ligand (52). We therefore next characterized co-localization of TLR2, TLR7 and MyD88 with phagosomes containing Bb in BMDMs. By confocal microscopy, we showed that in WT BMDMs there is colocalization of MyD88 (**Figure 3A**), TLR2 (**Figure 3B**) and and TLR7 (**Figure 3C**) with Bb-containing phagosomes. Signals from MyD88 and TLR2 distinctly overlay with Bb GFP signals from phagosomes showing evidence of coiled or degraded spirochetes (**Figure 3A-B, graphs**), but the intensity of MyD88 or TLR2 signal observed was higher with phagosomes containing degraded spirochetes. We also noted that TLR2 was expressed on the cell membrane and showed colocalization with surface-bound spirochetes (**Figure 3B**). TLR7 only showed strong signal with phagosomes containing partially degraded Bb but did not colocalize with surface-bound or recently internalized spirochetes (**Figure 3C**). In MyD88−/− BMDMs there is still colocalization of TLR2 with the phagosome (data not shown), but any TLR2 recruitment to the phagosome is not associated with pro-inflammatory signaling due to the lack of functional MyD88 protein (53). Taken together, these data confirm that endosomal TLR2, TLR7 and MyD88 colocalize to Bb-containing phagosomes to facilitate recognition of bacterial ligands and early response to infection.

**Figure 3:**
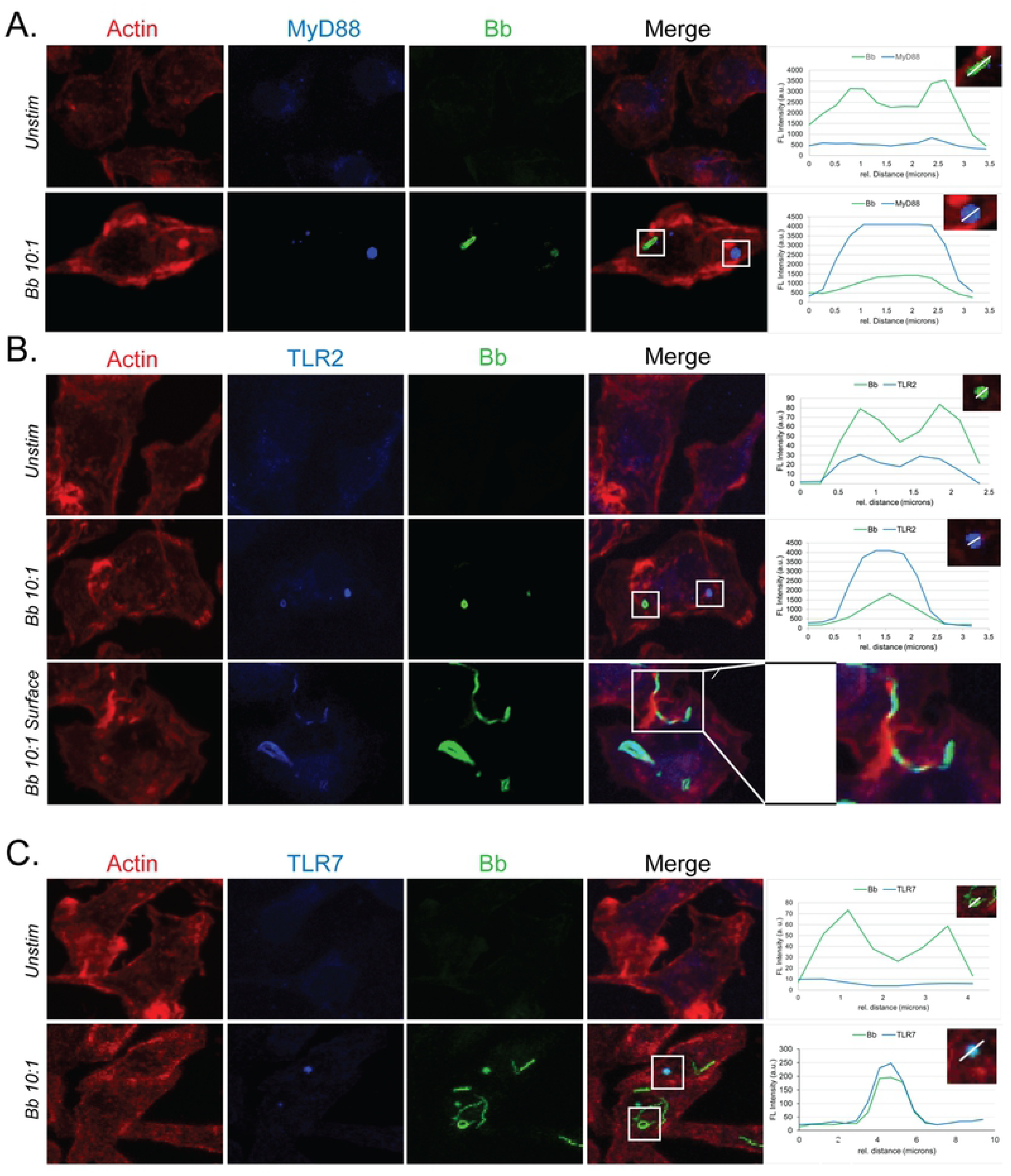
MyD88, TLR2 and TLR7 colocalize with Bb in phagosomes. (A-C) Confocal images and colocalization analysis of internalized Bb with MyD88 (A), TLR2 (B) or TLR7 (C) in WT BMDMs after stimulation at MOI 10:1. White box indicates phagosome depicted in inset. Large inset in (B) shows coiling pseudopod formation around Bb on cell surface. Graph shows the intensity of each indicated pixel marker across the white line (distance on x-axis). Green is Bb, blue is MyD88 (A), TLR2 (B) or TLR7 (7), and red is actin.

### Lack of MyD88 does not affect degradation of Bb in the phagosome

Degradation of the spirochete in the phagosome is crucial to expose bacterial ligands for recognition by endosomal TLRs (17). This process, known as phagosome maturation, requires reduction of phagosome pH and fusion with lysosomes (54). Given that both WT and MyD88−/− BMDMs bind and internalize Bb, we next sought to determine if spirochetes are similarly degraded in phagosomes with and without MyD88. Confocal images taken after a 6-hour stimulation at MOI 10:1 showed that both WT and MyD88−/− BMDMs contained degraded GFP+ Bb within the cell actin matrix (**Figure 4A and 4B**). To assess phagosome maturation, we quantitated recruitment of LAMP-1 to Bb-containing phagosomes by looking at colocalization of LAMP-1 and GFP fluorescence intensity (55). Both WT and MyD88−/− BMDMs showed comparable LAMP-1 and Bb colocalization in phagosomes (**Figure 4A and 4B**, graphs). Colocalization between Bb and LAMP-1 was measured in multiple phagosomes in BMDMs from both genotypes and no significant differences were found (**Figure 4C**). To confirm MyD88 signaling in response to Bb we also measured TNFα secretion after 1, 4 and 6 hours of incubation with spirochetes. WT BMDMs showed significant increase in TNFα secretion in the presence of spirochetes, while MyD88−/− BMDMs did not make TNFα protein (**Figure 4D**). MyD88−/− BMDMs also did not secrete IL-6, IL-12 or IL-10 (**Figure S3A-C**). Consistent with prior studies by Behera et al (2006), both WT and MyD88−/− BMDMs secrete the macrophage chemokine CCL2 (**Figure S3D).**

**Figure 4:**
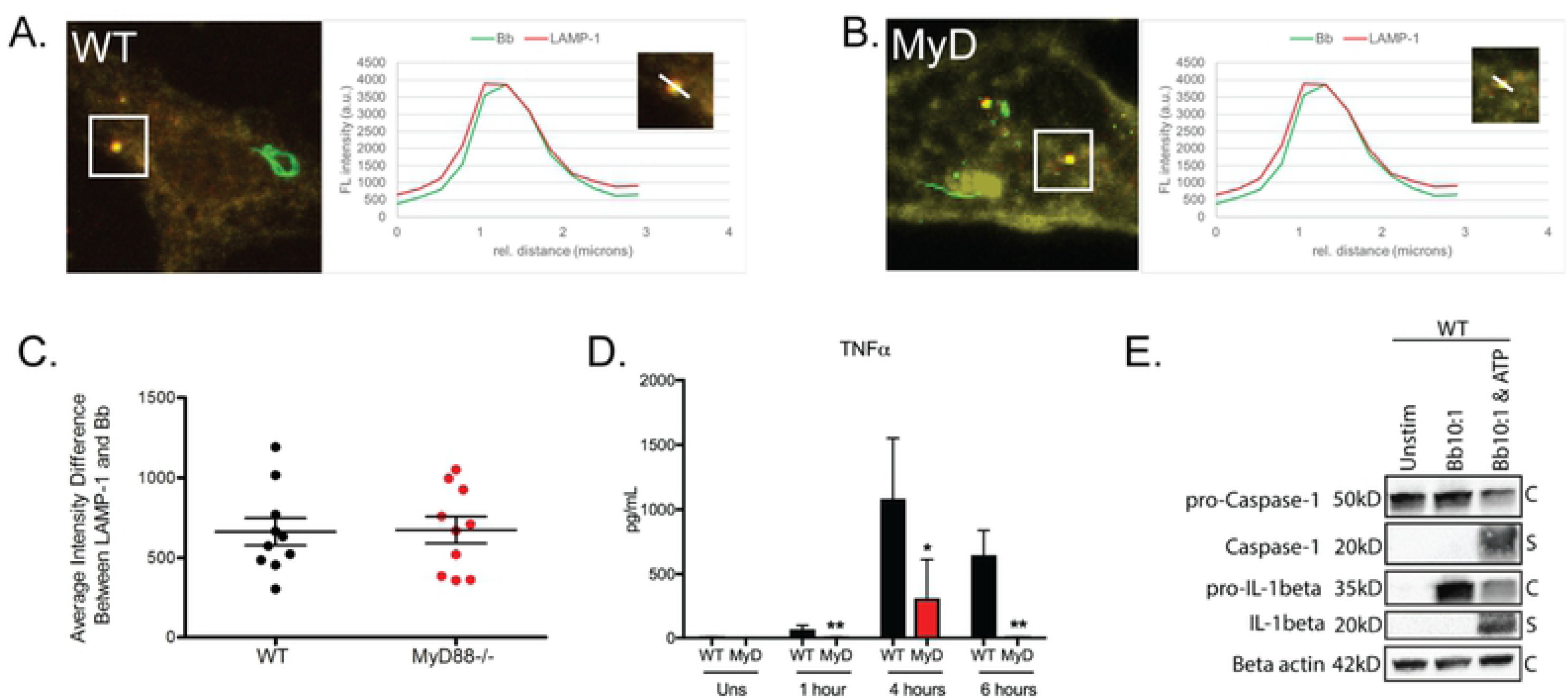
Colocalization of phagosome markers with internalized Bb in WT and MyD88−/− BMDMs. (A-B) Confocal images of WT (A) and MyD88−/− (B) BMDMs after 6 hours of stimulation with Bb at MOI 10:1, depicting colocalization of Bb-containing phagosomes with LAMP-1. White box indicates phagosome depicted in inset. Graph shows the intensity of each indicated pixel across the white line (distance on x-axis). Green is Bb, red is LAMP-1 and yellow is actin. (C) Quantitation of colocalization between Bb and LAMP-1 in 10 phagosomes of WT (black dots) and MyD88−/− (red dots) BMDMs by measuring intensity difference between LAMP-1 staining and Bb staining. (D) TNFα production in WT (black bars) or MyD88−/− (red bars) BMDMs after 1, 4 or 6 hours of stimulation at MOI 10:1. (E) Western blot of protein lysate isolated from WT BMDMs after 6-hour stimulation with Bb +/− ATP (C=cell lysate, S=supernatant). *p-value<0.05, **p- value<0.01, ***p-value<0.001, NS=not significant

### Bb ligand recognition appears to occur solely from within the phagosome

To test for the presence of bacterial products in the cytosol, we measured cleaved caspase-1, which is indicative of inflammasome activation. Western blot analysis of WT BMDM cell lysates and supernatants showed no activation of caspase-1 by stimulation of Bb alone (**Figure 4F**), which is consistent with previously published studies (56). However, in discordance with previous studies (57), we did not see cleavage of IL-1β (**Figure 4F**) unless exogenous ATP was added to the stimulation. To further confirm lack of NLRP3 inflammasome activation, we assessed <u>A</u>poptosis-associated <u>s</u>peck-like protein containing a <u>C</u>ARD (ASC) in BMDMs stimulated with either Bb or *Staphylococcus aureus* (Sa) for 30 minutes or 6 hours (**Figure S4**). As previously reported (58) (**Figure S4A and S4C**), ASC activation was observed with Sa, but no ASC was observed in BMDMs stimulated with Bb at 30 minutes or 6 hours (**Figure S4B and S4D**). Thus, recognition of Bb ligands appears to occur solely within the phagosome.

### MyD88-dependent signaling causes differential expression of genes in macrophages that promote the inflammatory response

Our results above emphasize the changes in inflammatory cell infiltrate and chemokine upregulation in tissues due to MyD88 signaling (**Figures 1 and S2**) and show that MyD88 expression in macrophages enhances their capacity to phagocytose spirochetes (**Figure 2**). To gain a better understanding of events that occur downstream of signaling by MyD88 which result in this phenotype presentation, we performed RNA sequencing on WT and MyD88−/− BMDMs stimulated with Bb for 6 hours (**Figure S5**). This time point was also selected based on our data in **Figure 4** showing comparable maturation in both WT and MyD88−/− BMDM phagosomes. We sequenced RNA from WT BMDMs at a MOI of 10:1 and MyD88−/− BMDMs at a MOI of 100:1 for a comparative analysis because the uptake percentages were not significantly different between the two cell phenotypes under these conditions (**Figure 5A**). Both WT and MyD88−/− BMDMs showed differentially expressed genes (DEGs) when compared to their respective unstimulated controls. We noted that the number of DEGs in WT BMDMs was much higher than in MyD88−/− BMDMs (2818 genes vs 141 genes respectively) (**Figure 5B**). We saw similar numbers of up- and down-regulated DEGs in WT BMDMs (**Figure 5B**). In the MyD88−/− BMDMs, approximately 83% of the DEGs were up-regulated (**Figure 5B**). We classified the DEGs into three categories for further analysis: genes differentially expressed only in WT BMDMs (MyD88-dependent); genes differentially expressed in both WT and MyD88−/− BMDMs (MyD88-independent); and genes that were differentially expressed only in MyD88−/− BMDMs (MyD88-privative) (**Figure 5C**).

**Figure 5:**
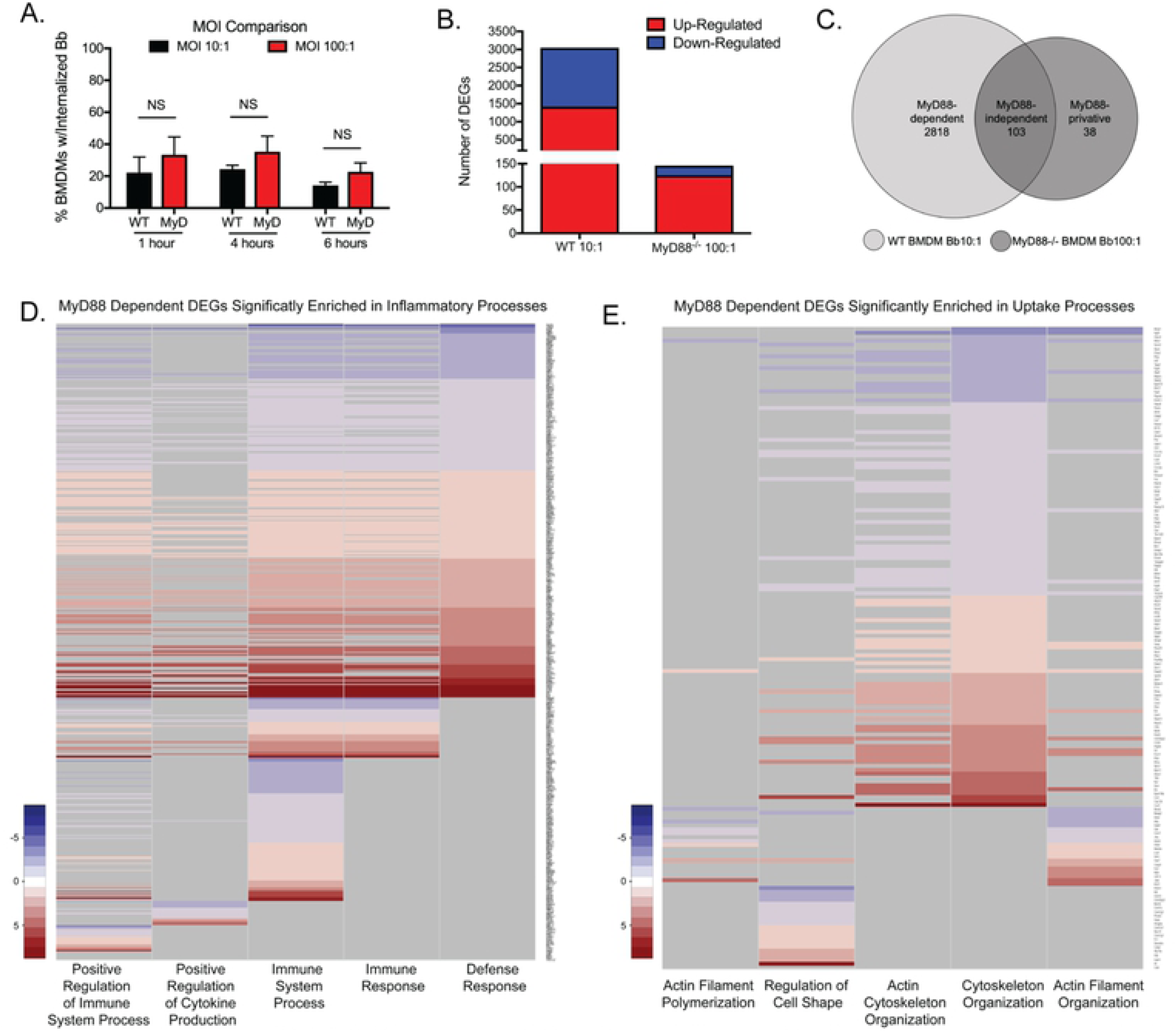
MyD88-dependent DEGs are significantly enriched in biological processes related to inflammation and uptake. (A) Comparison of Bb internalization by WT BMDMs (black bars) at MOI 10:1 with MyD88−/− BMDMs (red bars) at MOI 100:1. (B) Number of differentially expressed genes (DEGs) in Bb-infected WT and MyD88−/− BMDMs determined by RNA-sequencing. Red bar indicates number of upregulated DEGs and blue bar indicates number of down-regulated DEGs. Bar height represents total number of DEGs in each condition. (C) Venn diagram depicting DEG classification. MyD88-dependent genes (light gray, left) are only differentially expressed in WT BMDMs. MyD88-independent genes (center) are expressed in both cell types. MyD88-privative genes (dark gray, right) are only differentially expressed in MyD88−/− BMDMs. (D and E) Heat maps depicting fold-change values of MyD88-dependent DEGs enriched in specific biological processes. Red indicates positive fold-change increase, blue indicates negative fold-change increase, gray indicates no enrichment to biological process. (D) MyD88-dependent DEGs that significantly enriched to 5 indicated biological processes relating to inflammation. (E) MyD88-dependent DEGs that significantly enriched to 5 indicated biological processes relating to uptake.

Using these data, we performed a Gene Ontology (GO) enrichment analysis of each group of DEGs (**Supplemental File 1**). Because we showed above that in macrophages MyD88 affects both the inflammatory response and uptake of spirochetes we focused on biological processes relating to inflammation and phagocytosis in the MyD88-dependent DEGs. Multiple inflammatory biological processes were enriched (see supplemental files) but we concentrated on five processes with high numbers of gene hits: (i) Positive Regulation of Immune System Process, (ii) Positive Regulation of Cytokine Production, (iii) Immune System Process, (iv) Immune Response and (v) Defense Response. The MyD88-dependent DEG GO analysis resulted in the DEGs being broadly characterized into two different subsets. The first is comprised of DEGs enriched in all or almost all five processes and includes classic inflammatory genes, such as *Il6, Il12b, Il1b, Il23a* and *Tlr9* (**Figure 5D**). The second is comprised of DEGs that were enriched in only one or two of the five biological processes (**Figure 5D**). Examples of these DEGs included *Prdm1* (Positive Regulation of Immune System Process), which is involved in suppression of IFN-β production (59) and *Il17ra* (Positive Regulation of Cytokine Production), which facilitates the differentiation of neutrophils in response to IL-17 (60). The MyD88-independent DEGs also significantly enriched to the same five biological processes in their GO analysis but had fewer numbers of hits for each process (**Supplemental File 4**). The GO analysis of the MyD88-privative genes also showed significant enrichment to four of these inflammatory processes; Positive Regulation of Cytokine Production was not enriched in the MyD88-privative DEGs (**Supplemental File 4**). Overall this analysis indicates that MyD88 signaling results in activation of many processes and cellular elements that affect inflammation.

### Multiple genes associated with biological processes involved with phagocytosis are MyD88-dependent

Based on our observation that the presence of MyD88 enhances phagocytosis (**Figure 2**), we also analyzed whether any of the MyD88-dependent DEGs enriched to biological processes related to uptake. We identified 164 MyD88-dependent DEGs that enriched to five different biological processes relating to phagocytosis (Actin Filament Polymerization, Regulation of Cell Shape, Actin Cytoskeleton Organization, Cytoskeleton Organization, and Actin Filament Organization) (**Figure 5E**). Of particular interest, *Daam1* and *Fmnl1*, encoding two proteins known to play a role in phagocytosis of Bb (15, 37), were differentially expressed in an MyD88-dependent manner (**Supplemental File 3**). Daam1, which was upregulated, is a formin protein that bundles actin fibers together to increase stability of coiling pseudopods, which are more adept at capturing the highly motile spirochetes (61). Fmnl1, which was down-regulated, is also a formin protein that severs actin branches to promote polymerization and increase filopodia protrusion (61).To better understand how these genes relate to other genes commonly associated with phagocytosis, and to understand which pathways are turned on or off, we first separated the 164 genes into two groups based on upregulation or downregulation. We then entered the list of genes from each group into the GeneXplain software to map interactions among the proteins they encode. Pathways including the uptake genes with up to 10 intermediates between them were computed. Because Daam1 and Fmnl1 have been previously implicated in phagocytosis of Bb, we then merged all pathways that included either *Daam1* or *Fmnl1* into protein interaction networks (**Figure 6**). The network mapping *Daam1* chains (**Figure 6A**) shows related pathways that are upregulated and includes *Rhoa, Akt1, Rac1* and *Cdc42* as genes that code for proteins which appear as intermediates on the network, meaning that their genes weren’t differentially expressed in our analysis. (**Figure 6A, red boxes**). *Daam1* regulates *Rhoa* activity, which controls *Cdc42, Rac1* and *Akt1*. *Cdc42* activates a Rho GTPase, *Rhoq*, which is upregulated in response to Bb. *Rac1* and *Akt1*, when translated, both activate multiple proteins whose corresponding genes are also upregulated, indicating that while the genes for these intermediate proteins aren’t differentially expressed, they are still active in macrophages that have been stimulated with Bb. In contrast to *Daam1, Fmnl1* was downregulated in response to Bb. The network mapping *Fmnl1* chains (**Figure 6B**) also highlights other pathways and genes that are downregulated. This network also includes the genes *Cdc42* and *Akt1* which code for intermediate proteins. However, the network shows that the genes which code for proteins that inactivate *Cdc42* and *Akt1* are being downregulated in an MyD88-dependent manner. The downregulation of these inhibitors indicates that the activity of these proteins is important in the macrophage response to Bb, consistent with the *Daam1* network above. Taken together, these data suggest that MyD88 signaling upregulates multiple gene products involved in regulating macrophage membrane protrusions. Upregulation of these genes likely contributes to the reorganization of cell machinery that enhances the capability of the WT macrophage to take up spirochetes.

**Figure 6:**
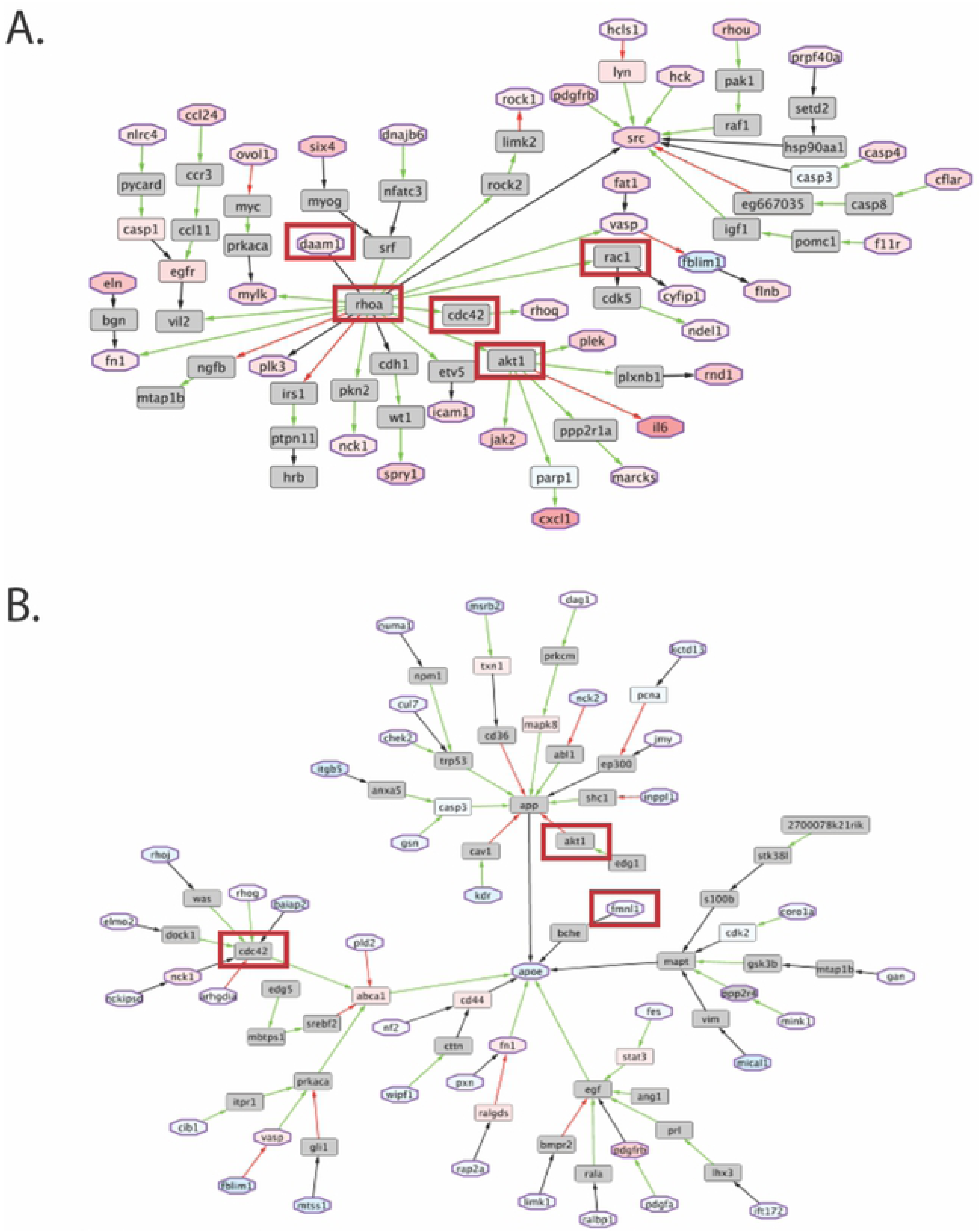
MyD88-dependent DEGs that significantly enrich to uptake processes show direct and indirect connections with key proteins involved in Bb phagocytosis. (A) Network of MyD88-dependent uptake DEGs that are upregulated. Octogonal nodes with purple borders indicate genes that significantly enriched to uptake biological processes. The varying degree of red or blue hue in select nodes correlates with the gene’s Log2 Fold Change value. Red indicates positive fold change and blue indicates negative fold change. Gray nodes represent genes that were not differentially expressed. Red boxes indicate genes of interest reviewed in the results. (B) Network of MyD88-dependent uptake DEGs that are down-regulated. The same parameters and color scale applied in (A) was used.

### Similar inflammatory and chemotactic processes are enriched regardless of MyD88-mediated signaling, but utilize different regulatory proteins

MyD88-dependent mechanisms of inflammation have been well characterized, but little work has been done to understand the drivers of Bb-induced inflammation in the absence of MyD88. To address this issue, we next completed a comprehensive analysis to determine how the DEGs are regulated within Bb-infected macrophages, both in the presence or absence of MyD88. We first identified transcription factors with potential binding sites in the promoter regions of the DEGs for each of the three subsets (66 for MyD88-dependent, 201 for MyD88-independent, and 39 for MyD88-privative). We then identified master regulator proteins upstream of these transcription factors and performed a GO enrichment analysis using the same parameters described in **Figure 5**. The enrichment analysis of MyD88-independent master regulators revealed processes involved in cell metabolism and homeostasis, which did not give much insight into differences in mechanisms of inflammation driven by the presence or absence of MyD88. Therefore, we focused on analysis of MyD88-dependent and MyD88-privative master regulators. In light of our findings that Bb-infected MyD88−/− mice showed increased macrophage and neutrophil infiltrate in heart tissue (see **Figure 1**), we investigated whether any master regulators enriched to inflammatory and/or chemotactic biological processes. Interestingly, similar biological processes significantly enriched in inflammation were both identified in the MyD88-dependent (including *MyD88, Irak2* and *Ly96*) and MyD88-privative (including *Vcam1* and *Cxcl2*) master regulators (**Figure 7A and 7C**), but the individual master regulators involved were different for each subset (**Figure 7A**). Importantly, over three times as many master regulators were identified for the MyD88-dependent DEGs than the MyD88-privative DEGs (**Figure 7A**), suggesting that MyD88 signaling controls activation of more master regulators in the cell to initiate DEGs (see **Figure 5**) and enables the cell to perform unique processes in response to Bb.

**Figure 7:**
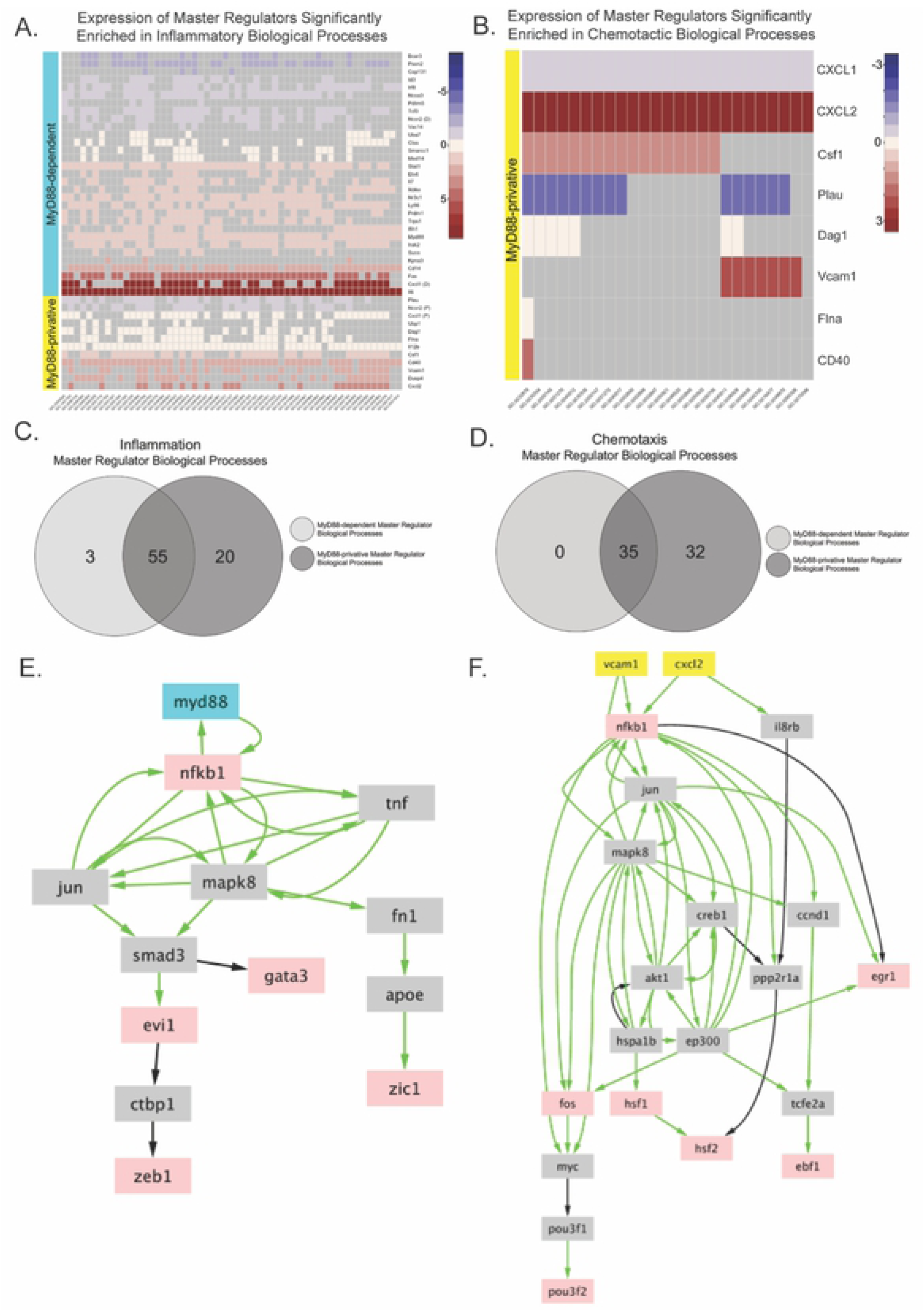
Master regulators of MyD88-dependent and MyD88-privative DEGs control similar inflammatory processes but different chemotactic processes. (A and Heat map showing fold change of master regulators enriched in inflammation (A) or chemotaxis (B) in WT (cyan) or MyD88−/− (yellow) BMDMs. GO numbers for significantly enriched BP are indicated on the x-axis. (C and D) Venn diagrams comparing biological processes (BP) relating to inflammation (C) or chemotaxis (D) significantly enriched between MyD88-dependent (light gray) and MyD88-privative (dark gray) master regulators. (E-F) Protein networks of significant master regulator proteins. Green arrows indicate positive regulation from one protein to another. Black arrows indicate unknown effect of one protein on another. Proteins in gray boxes represent intermediates and red boxes indicate transcription factors. (E) Protein network of MyD88 (blue box) and downstream intermediates. (F) Protein network of MyD88-privative master regulators (yellow boxes) that significantly enriched to chemotaxis-related BP.

### MyD88 is a master regulator for transcription factors that control the MyD88-dependent DEGs enriched in uptake processes

To better understand the phenotypic differences related to phagocytosis between the two genotypes (see **Figure 2)**, we identified transcription factors that map to promoter regions of DEGs related to phagocytosis. We then used OCSANA in Cytoscape to link MyD88 (as a master regulator) with transcription factors that map to DEGs in this specific subset. Of these transcription factors, three were expressed exclusively in WT BMDMs; *Zic1, Zeb1* and *Gata3*. Based on this information we constructed a network illustrating potential links between MyD88-mediated signaling and enhanced phagocytic capability seen in WT cells (**Figure 7E**). Analysis of these transcription factors revealed that Zic1, Zeb1 and Gata3 have the capacity to bind to the promoter regions of several of the MyD88-dependent DEGs that significantly enriched to processes associated with bacterial uptake. Zic1 is controlled by the intermediate protein ApoE, which is known to play a role in cholesterol metabolism in macrophages (62) and the absence of ApoE increases Bb burdens in experimentally infected mice (62).

### MyD88-privative master regulators are involved in multiple chemotactic biological processes not enriched in WT BMDMs

We also observed significant overlap between the chemotactic biological processes enriched in MyD88-dependent and MyD88-privative master regulators. However, MyD88-privative master regulators significantly enriched to multiple biological processes involved in chemotaxis that were not enriched in MyD88-dependent master regulators (**Figure 7B and 7D**), suggesting that the lack of MyD88 signaling allows for increased up-regulation of processes to facilitate cell migration into the tissues. The MyD88-privative master regulators involved in these chemotactic processes also enriched to inflammatory processes (**Figure 7**), suggesting that Bb may trigger other signaling cascades which induce inflammation more skewed to cell recruitment and localization. We then generated a network of two MyD88-privative master regulators that were up-regulated and significantly enriched to multiple chemotactic biological processes (**Figure 7F**). Interestingly, *Cxcl2* was differentially expressed in the absence of MyD88 and also mapped as a master regulator in the network. Most importantly, both networks shown in **Figure 7E and 7F** provide potential targets for knock-out studies to test if they contribute to inflammation in humans.

### The absence of MyD88 drives unique chemokine transcript expression profiles in heart tissue during Bb infection

The analysis of RNA-sequencing data revealed that expression of several chemokines was MyD88-independent. To determine whether these chemokines were expressed in the absence of MyD88 *in vivo*, we performed RT-PCR of the chemokine genes from total RNA isolated from heart and patellofemoral joint tissues from Bb-infected mice (*Ccl2, Ccl9, Cxcl2, Cxcl3, Cxcl10*). *Ccl2* and *Ccl9* are macrophage-specific chemokines (63). Significant transcriptional differences between WT and MyD88−/− mice were seen 14 and 28 DPI (**Figure S6**). WT hearts showed marked upregulation of all chemokines tested at 14 DPI in comparison to uninfected WT controls, with the exception of *Ccl2* (**Figure S6A**). In contrast to WT mice, *Ccl9, Cxcl2, Cxcl3 and Cxcl10* were down-regulated in MyD88−/− infected hearts at 14 DPI compared to uninfected controls (**Figure S6**). However, at 28 DPI *Ccl9* was upregulated in MyD88−/− heart tissue, but down-regulated in WT hearts (**Figure S6C**). In joint tissue MyD88−/− mice did not show any significant upregulation of chemokines 14 DPI, but *Ccl2, Cxcl2, Cxcl3* and *Cxcl10* were up-regulated 28 DPI (**Figure S6)** These chemokine transcripts were downregulated in WT mice 28 DPI. Interestingly, *Cxcl2* and *Cxcl10* were not significantly upregulated in heart tissue at the same time point (**Figure S6**). These chemokines have high affinity for the IL-8 receptor (64), and may facilitate recruitment of other inflammatory cells (other than macrophages) to joint tissue in the absence of MyD88. Overall, these data suggest that in the absence of MyD88, cells in heart tissue show different inflammatory gene expression in response to Bb, with increased production of select chemokines that can facilitate macrophage recruitment.

## DISCUSSION

Previous studies by our group have emphasized that uptake and degradation of Bb by phagocytic cells, including monocytes and macrophages, are critical in eliciting the inflammatory response to the bacterium (14, 17-19, 29). The findings from these studies, as well as others (28), show that the adaptor protein MyD88 plays a critical role in bacterial uptake and phagosomal signaling in macrophages. In the current study, we provide further evidence that the macrophage is a key driver of inflammation, even in the absence of MyD88. We also show that while MyD88 has a significant impact on spirochetal uptake, phagosome maturation and bacterial degradation are not affected. Of particular novelty, phagosomal signaling cascades induced by Bb ligands in macrophages trigger a number of inflammatory and chemotactic pathways. Moreover, the inflammatory processes are mediated by different regulatory proteins depending on whether MyD88 is present or absent, while induction of several chemotactic processes occurs independently of MyD88. In-depth analysis of these signaling cascades allowed us to identify previously underappreciated MyD88-dependent master regulators and transcription factors which can lead to enhanced spirochetal uptake and clearance.

The importance of MyD88 signaling in Bb clearance from infected murine tissues is well recognized (40-42), but the tissue specific kinetics of spirochetal clearance have not been fully defined. In the current study, we quantified Bb burdens in multiple tissues at early (14 DPI), intermediate (28 DPI) and late (56 DPI) time points post-infection to more fully establish bacterial clearance kinetics and how the rate of clearance is affected by the presence of MyD88. As in these prior studies, our results (see **Figure S1)** confirm that MyD88 signaling in Bb-infected mice affects spirochete clearance. In contrast to prior work (42), we did not observe significant differences in spirochete burdens at 14 DPI. In bladder, skin, ear, and joint tissue the lack of MyD88 results in persistent Bb burdens that remain elevated even 56 DPI. Given that there was a reduction in Bb burdens in MyD88−/− mouse hearts 56 DPI, we postulate that MyD88 signaling in macrophages is a necessity for early control of Bb burdens in heart tissue, The significant decrease in Bb burdens in MyD88−/− murine heart tissue by 56 DPI, but not in other tissues taken from the same mouse, indicates that tissue-specific inflammatory responses dependent on MyD88 signaling can impact spirochete clearance.

Bb-induced tissue-specific differences in inflammation severity and types of cell infiltrate in the mouse model of infection have been reported in prior studies (5, 6, 40, 41, 44, 65, 66). Inflammation in mouse joint tissue is more neutrophilic (40), while heart inflammation is more macrophage-driven (10). In this study we focused on characterization of inflammatory macrophages because of their role in early recognition and clearance. Interestingly, our studies show that heart tissue in MyD88−/− mice had significantly more inflammation and macrophage infiltrate 28 DPI than Bb-infected WT hearts (**Figure 1**). These findings suggest that MyD88 is not required for immune cell recruitment since macrophage-specific chemokine transcript is significantly upregulated and macrophages readily appear in infected heart tissue despite the lack of MyD88 signaling. Whether MyD88 signaling affects other factors necessary for macrophage recruitment, such as integrin expression, has not been studied. Lack of β2/CD18 integrin expression increases inflammation severity (11), and lack of CD11a increases bacterial burdens in the heart (67, 68). Our results also suggest that macrophages are not the only cell driving spirochete clearance *in vivo*. Infected MyD88−/− mouse hearts showed significant neutrophil infiltration 28 DPI, greater than WT mice, suggesting that, similar to joints, neutrophils also play a role in chemokine production and Bb clearance in the heart (69, 70). Whether MyD88 signaling affects the ability of neutrophils to efficiently capture spirochetes is not known. Taken together, our *in vivo* Bb infection studies reveal that, in heart tissue, the absence of MyD88 signaling results in recruitment of more phagocytic cells (i.e. macrophages and neutrophils) and suggest that these cells still play an important role in controlling bacterial loads despite their diminished ability to take up spirochetes.

To better understand the contribution of MyD88 to spirochete binding, uptake, degradation and signaling by macrophages, we transitioned to an *ex vivo* model. Murine macrophages lacking MyD88 show a phagocytic defect when stimulated with Bb *ex vivo* compared to WT macrophages. This defect in uptake has been previously demonstrated in macrophage stimulation experiments with other bacteria strains (31, 34, 35, 38). Our results reveal that binding of Bb is not affected and that this phagocytic defect is not dependent on length of stimulation (i.e. time dependent) and is only slightly rescued by increasing the MOI. Thus, in the absence of MyD88, macrophages are still capable of binding and taking up the LD spirochete, but MyD88 signaling enhances the efficiency of Bb phagocytosis by macrophages. We also show here that when stimulated *ex vivo*, the macrophage response to Bb is driven by the signaling cascades induced by MyD88 as a result of bacterial ligands engaging TLR2 and TLR7 receptors in the phagosome. Recognition in the phagosome is driven by degradation of bacteria, since more TLR2, TLR7 and MyD88 marker intensity were observed colocalizing with degraded spirochetes. Our results in **Figure 4** indicate that in the context of Bb infection, MyD88 is not required for phagosome maturation, evidenced by the recruitment of LAMP-1 to Bb-containing phagosomes in MyD88−/− BMDMs. This is in contrast to Blander *et al.*, who published that MyD88−/− BMDMs infected with *S. aureus* or *E. coli* did not colocalize with either Lysotracker or LAMP-1 to the same degree as WT BMDMs (38). One possible explanation for the different findings with our study is that the recruitment of LAMP-1 is delayed in MyD88−/− BMDMs, given that Blander *et al.* measured phagosome maturation at an earlier time point than in our studies. This explanation is supported by Yates *et al*. who showed slightly delayed acidification in MyD88−/− BMDMs stimulated with TLR2 or TLR4 ligands for 40 minutes (46). The fragility of the Bb membranes also suggests that perhaps less acidification of the phagosome is needed to expose Bb PAMPs.

Reduced uptake of bacteria in macrophages lacking MyD88 is a phenotypic trait that has been extensively detailed (28, 31-35), but not well-understood. To better understand the relationship between MyD88 signaling and phagocytosis, we used a computational systems biology approach. In prior studies, addition of TLR3 ligands to Bb stimulation of MyD88−/− BMDMs significantly rescues uptake (36), suggesting that in the absence of MyD88 TRIF signaling can activate pathways that result in similar actin rearrangement in the cell. This signaling was shown to be mediated through PI3K (36), but interestingly PI3K was not differentially expressed in our macrophage stimulation. However, our network analysis from RNA-sequencing data identified DEGs that are upregulated downstream of common phagocytosis effector proteins. *Rhoq*, activated by *Cdc42* codes for TC10, a protein involved in generating long filopodia protrusions (71). The gene *Cyfip1*, which encodes a part of the WAVE complex that regulates actin polymerization (72), was also upregulated according to the network through *Rac1* protein interactions. The WAVE complex has higher involvement with lamellipodia formations (73). It is likely that MyD88 controls transcription factors that upregulate these genes to promote phagocytosis through formation of coiling pseudopods, which are more similar to lamellipodia, rather than through straight filopodia protrusions. In addition to MyD88, studies indicating that TLR2 can utilize TRIF have also been completed, but this interaction only appears to contribute to the inflammatory response rather than spirochete uptake (18, 74). More recently, the leukotriene LTB_4_ has been shown to promote phagocytosis of Bb by macrophages (75), but in our BMDM sequencing data we did not find differential expression of *Ltb4* or its receptor *Ltb4r1*. This could possibly be due to the later time point we selected for sequencing. It has also been shown that spleen tyrosine kinase (Syk) has an important role in phagocytosis of Bb via integrin binding (76). The Syk gene (*Syk*) is significantly upregulated in an MyD88-dependent manner, suggesting that MyD88 drives over-expression of *Syk* to increase phosphorylation and activation of proteins involved in generating actin branches. However, in our GO analysis *Syk* was not one of the 164 MyD88-dependent genes that enriched to uptake biological processes, and the transcription factor Zic1, which has binding sites in the promoter regions of a significant number of these genes, is not predicted to bind in the promoter region of *Syk*. Zic1 was of particular interest to us because it appeared downstream of MyD88 in our network analysis (**Figure 6**) and is controlled by the intermediate protein ApoE. Mice lacking ApoE have increased bacterial burdens when infected with Bb (62), suggesting that ApoE signaling plays a role in cell remodeling processes necessary to enhance uptake. In addition, a link between Bb phagocytosis and cholesterol has been postulated by Hawley *et al*. who showed that CR3, a known phagocytic receptor for Bb, is recruited to lipid rafts with the co-receptor CD14 (77). Thus, it is possible that MyD88 upregulates ApoE to enhance lipid rafts on the macrophage membrane, which can potentiate signaling to enhance uptake and provide scaffolding for proteins involved in actin remodeling.

Our computational analysis also supported that there are non-canonical sources of inflammation in MyD88−/− mice. Our results suggest that there is possibly another receptor recruited to the phagosome that initiates chemokine production upon recognition of a Bb ligand (see **Figure S6**). Another mechanism for triggering chemokine production may be that the TLR receptors are utilizing another adaptor protein to transmit signals out of the phagosome, as postulated by Petnicki-Ocwieja *et al.* (74). The production of chemokines facilitates recruitment of cells to spirochetes infected tissues, but the increased cell recruitment in MyD88−/− mice, coupled with higher Bb burdens suggests that the infiltrating immune cells are not efficient at clearing spirochetes. We hypothesize that in order to compensate for their lack of efficiency at uptake, more cells need to be recruited. This progression of events may not be true for joint compared to heart tissue, since our results show that the chemokine responses are different in each tissue. Network analysis of DEGs from macrophages identified multiple master regulators that could be controlling production of these chemokines, but further work needs to be done to determine if these master regulators are in fact active in macrophages containing Bb. Additional studies to test whether acidification of the phagosome is required for Bb-induced chemokine production will also give insight into which ligand-receptor interaction induces this response.

In summary, our results emphasize that the macrophage has a very important role in both recognition and clearance of Bb and is at the epicenter of the immunologic response to spirochete infection, particularly in relation to Lyme carditis. The findings from these studies have also advanced our understanding of how phagosomal signaling drives spirochete uptake, recognition and inflammation. The adaptor protein MyD88 plays a critical role in these processes. Initial phagocytosis of Bb by macrophages does not require MyD88, but once taken up, recognition of Bb ligands exposed upon spirochete degradation occurs through endosomal TLRs which trigger MyD88-mediated signaling cascades. This signaling results in cell remodeling to enhance phagocytosis, as indicated by our *ex vivo* data, and allows macrophages to more efficiently internalize and clear the highly motile spirochetes by using more dynamic membrane protrusions. Further studies using the targets identified in these experiments may also provide insight into understanding the importance of phagocytosis in other bacterial infections. We can use similar techniques to look at the role of the macrophage response and MyD88 signaling in human macrophages, with the goal of increasing our understanding of the clinical spectrum associated with Lyme disease pathogenesis.

## MATERIALS AND METHODS

### Mice

Female 6-8-week-old C57BL/6J wild type (WT) and C57BL/6J MyD88−/− (MyD88−/−) mice used in these studies were obtained from breeding colonies maintained in the UConn Health (UCH) Center for Comparative Medicine facility according to guidelines set by the UCH Institutional Animal Care and Use Committee. Original WT breeding pairs were purchased from The Jackson Laboratory (Bar Harbor, Maine). Original MyD88−/− breeding pairs were kindly provided by Dr. Egil Lien at the University of Massachusetts with permission from Dr. S. Akira in Osaka, Japan. Disruption of the murine MyD88 gene was confirmed through PCR (53). Both WT and MyD88−/− breeding colonies are maintained on the antibiotic Sulfatrim (sulfomethoxazole [40 mg/mL] + trimethoprim [8 mg/mL]) diluted in water 1:50, which has been previously shown to not impact the degree of Bb infection (42).

### Ethics Statement

All experiments in this work involving the use of animals or isolation of primary cells from animals were monitored and approved by the UConn Health Institutional Animal Care and Use Committee under protocol #101388-0819. The UConn Health Institutional Animal Care and Use Committee is accredited by The Association for Assessment and Accreditation of Laboratory Animal Care (AAALAC).

### Bacterial Strains

Low-passage virulent wild-type strain 297 (78) or a strain 297 isolate containing a stably-inserted copy of green fluorescent protein (GFP) under the control of the constitutively-expressed *flaB* promoter (Bb914) (79) were maintained in Barbour-Stonner-Kelly (BSK)-II media supplemented with normal rabbit serum and gentamicin (50 μg/μl) (79). Cultures were grown at 23°C for at least one week prior to being shifted to 37°C as previously described (79). Spirochetes were centrifuged at 3300 × *g* for 20 minutes at 4°C and resuspended in either BSK-II for *in vivo* experiments or DMEM (Gibco, 15630-080) supplemented with sodium pyruvate (Gibco, 11360-070) and HEPES (Gibco, 15630-080) for *ex vivo* experiments. After resuspension, the cultures were counted by dark-field microscopy using a Petroff-Hausser counting chamber (Hausser Scientific) and diluted accordingly. *Staphylococcus aureus* (Sa) was cultured and fluorescently labeled with FITC as previous described (80).

### Mouse Infection

Sex- and age-matched WT and MyD88−/− mice were used in every experiment. Mice were injected intra-dermally on their backs with either 1 × 10^5^ temperature-shifted Bb diluted in 50 μL BSK-II or sham-inoculated with an equal volume of media alone and then sacrificed 14-, 28- or 58-days post-infection. Infection was confirmed at 2 weeks post-infection by serology and culturing of ear tissue in BSK-II. Cultures were examined weekly for spirochetes by darkfield microscopy for at least 4 weeks. Serum was tested against Bb protein lysates via chemiluminescent immunoblot using goat anti-mouse HRP-conjugated IgG (GE NA931) as previously described (81).

### Quantitation of Bb Burdens in Mouse Tissues

Mouse heart, tibiotarsal joint, patellofemoral joint, bladder, abdominal skin, and ear tissues were harvested at the time of sacrifice. Tissues were digested overnight in 0.1% collagenase Type I in PBS (Gibco, 17100-017) at 37°C, then further digested with 0.2 mg/mL proteinase K solution (200 mM NaCl, 20 mM Tris-HCl pH 8.0, 50 mM EDTA, 1% SDS, 0.2 mg/mL proteinase K (Machery-Nagel, 740506) in dH_2_O) overnight at 56°C. DNA from a small volume of digested tissue was isolated using a genomic DNA purification kit (Machery-Nagel, 740952) according to manufacturer’s instructions. Spirochete burdens were quantitated by qPCR using TaqMan-based assays for Bb *flaB* and murine nidogen (*nido*). The primers and probe used to detect *flaB* were flaB-F (5-CTTTTCTCTGGTGAGGGAGCTC-3’) and flaB-R (5’-GCTCCTTCCTGTTGAACACCC-3’) and flaB-probe (5’-FAM-CTTGAACCGGTGCAGCCTGAGCA-3’-BHQ1). The primers and probe used for *nidogen* were nidogen-F (5’-CCCCAGCCACAGAATACCAT-3’), nidogen-R (5’-AAAGGCGCTACTGAGCCGA-3’), and nidogen-probe (5’-FAM-CCGGAACCTTCCCACCCAGC-3’-BHQ1). The copy numbers for *flaB* and *nidogen* genes were calculated by the Bio-Rad software (version 3.1) based on standard curves (10^7^ – 10^2^ copies) generated from cloned versions of the corresponding amplicon (82, 83). *flaB* copy numbers were normalized to copies of *nidogen*. General statistical analysis was performed using GraphPad Prism 4.0 (GraphPad Software, San Diego, CA), using an unpaired Student *t* test. For each experiment, both the standard deviation and the standard error of the mean were calculated. *p* values of <0.05 were considered significant.

### Histopathology

Mouse patellofemoral joints and hearts were fixed for one week in 10% buffered formalin. Joints were decalcified for 48 hours after fixation. Tissues were then embedded in paraffin and 5 μm sections were floated onto glass slides using a 40°C water bath. Tissues were stained in hematoxylin solution for 90 seconds and Eosin Y solution for 18 seconds. Joints were scored on a scale of increasing severity from 1-3 by a pathologist in a blinded fashion for arthritis severity. Hearts were scored on a scale of increasing severity from 1-3 by a pathologist in a blinded fashion according to type and amount of cell infiltrate. General statistical analysis was performed using GraphPad Prism 4.0 (GraphPad Software, San Diego, CA), using an unpaired Student *t* test. For each experiment, both the standard deviation and the standard error of the mean were calculated. *p* values of <0.05 were considered significant.

### qRT-PCR Analysis of Infected Mouse Tissues

Mouse patellofemoral joints and hearts were homogenized using an electric homogenizer with toothed blades in tissue lysis buffer (Machery-Nagel, 740962) containing beta-mercaptoethanol (1:100) (Bio-Rad, 161-0710) and Triton-X-100 (1:100) (Fisher, BP151-100). RNA was isolated from tissue lysates using a Nucleospin RNA Kit (Midi column, Machery-Nagel, 740962) according to the manufacturer’s fibrous tissue protocol. RNA was quantified using a Nanodrop spectrometer (Thermo-Fisher). cDNA was generated from 2.5 nanograms total RNA using a high capacity cDNA RT kit (Applied Biosystems, 4368814) according to manufacturer’s instructions. Gene-specific qRT-PCR analyses were performed as previously described (29). Expression levels for *Aif1* (Mm00479862_g1), *Casp1* (Mm00438023_m1), *Ccl2* (Mm00441242_m1), *Ccl9* (Mm00441260_m1), *Cxcl10* (Mm99999072_m1), *Cxcl2* (Mm00436450_m1), *Cxcl3* (Mm01701838_m1), *Ifnb* (Mm00439546_s1), *Il12a* (Mm00434169_m1), *Il1b* (Mm01336189_m1), *Il6* (Mm00446190_m1), *Itgam* (Mm00434455_m1), *Mmp9* (Mm00442991_m1), *Ncf2* (Mm00726636_s1), *Nlrp3* (Mm00840904_m1), *Nod2* (Mm00467543_m1), and *Tnfa* (Mm00443258_m1) were normalized to *B2m* (Mm00437764_m1) and the fold changes in gene expression relative to the unstimulated or uninfected control were calculated using the 2^-ΔΔCt^ method (84).

### BMDM Stimulation

Bone marrow-derived macrophages (BMDMs) were isolated from 6-8-week-old WT and MyD88−/− mice as described previously (18). Single cell macrophage suspensions were seeded into either 12-well tissue culture-treated plates at a concentration of 1 × 10^6^ cells/ml per well or 1 × 10^5^ cells/500 μL per well in 8-chamber cell microscopy slides. Slides and plates were then incubated overnight at 37°C/5% CO_2_ to allow cell adherence before experimentation. Cells were incubated for either 0.5, 1, 4 or 6 hours at 37°C/5% CO_2_ with live GFP-Bb or labeled Sa at multiplicities of infection (MOIs) of either 10 or 100. Stimulation media was DMEM supplemented with 1% sodium pyruvate and 1% HEPES. For stimulations looking at the effect of Bb uptake, Cytochalasin D (Sigma, C8273-1MG) was added to wells at a concentration of 10 μg/mL and incubated at 37°C for 1 hour before Bb. At the end of the incubation period, culture supernatants were collected and stored at −80°C until cytokine analysis. Cells stimulated in chamber slides were processed for confocal microscopy. Cells stimulated in 12-well plates were processed for RNA extraction. All culture media and reagents were confirmed free of LPS contamination (<10 pg/ml) by Limulus amoebocyte lysate assay quantification (Cambrex, MA).

### Confocal Microscopy

After stimulation, BMDMs were fixed in 2% paraformaldehyde with 0.05% Triton-X-100 (Fisher, BP151-100) for 10 minutes. Slide wells were then incubated with 5% bovine serum albumin (BSA) solution in PBS overnight at 4°C to block non-specific antibody binding. The next day, cells were stained with different combinations of anti-GFP (Thermo Scientific A-21311, 1:100), phalloidin conjugated with Alexa Fluor 647 (Biolegend 424205, 1:20), anti-MyD88 (Santa Cruz 11356, 1:100), anti-TLR2 (eBioscience 14-9021-82, 1:100), anti-TLR7 (R&D MAB7156, 1:100), anti-ASC (Santa Cruz 22514-R, 1:100) and anti-LAMP-1 (eBioscience 14-1071-82, 1:100). A secondary antibody was used to detect anti-MyD88, anti-TLR2, anti-ASC and anti-LAMP-1 (Molecular Probes Alexa Fluor 568, A11036) (1:200). Incubations with primary and secondary antibodies were done for 1 hour each at room temperature; slide wells were washed following each incubation three times with PBS supplemented with 0.5% Tween-20, with a final wash in distilled H_2_O before mounting. After antibody staining, slides were mounted using Vectashield (Vector H-1000) and imaged using a Zeiss 880 confocal microscope. Image processing and analysis were performed using ImageJ (NIH, v1.41b). Colocalization values were determined by first analyzing profile plots in ImageJ (Plug-in: “Plot Profile”) across ten different phagosomes for each cell genotype and then calculating the average difference between the fluorescence intensity curves of the markers of interest (i.e., LAMP-1 and Bb). Binding percentages were calculated by imaging 100-200 cells using a confocal microscope and then measuring the ratio of cells containing at least one surface-bound or internalized spirochete to the total number of cells imaged for each condition, represented as %BMDMs interacting w/Bb. Uptake percentages were calculated by imaging 100-200 cells using a confocal microscope and then measuring the ratio of cells containing at least one internalized spirochete to the total number of cells imaged for each condition, represented as %BMDMs w/internalized Bb.

### Western Blotting of BMDM Supernatants and Lysates

Protein lysates were generated from BMDM cell culture lysates and supernatants after Bb stimulation. In these experiments, adenosine triphosphate (ATP) (Sigma, 3A6419-1G) was added to WT BMDMs (already stimulated with Bb for 5 hours) 1 hour prior to harvest for generation of lysates. Supernatants were treated with an equal volume of methanol and ¼ volume of chloroform, vortexed and spun at 16000 × *g* for 10 minutes. After removal of the upper phase, 500 μL of methanol was added to the intermediate phase, which was then vortexed and spun at 16000 × *g* for 10 minutes. The pellets were then dried at room temperature, resuspended in 30 μL of 2x Laemmli buffer and incubated in a 37°C water bath until proteins became soluble. BMDMs were lysed using RIPA buffer at −80°C and spun at maximum speed for 10 minutes. Protein pellets were resuspended in 2x Laemmli buffer. Lysates were boiled at 99°C for 10 minutes and run on a 12.5% SDS-PAGE gel at 140V for 1 hour (5μL per lane, 15 lanes). Proteins were then transferred to nitrocellulose membranes (Bio-Rad 162-0177) at 20V for 20 minutes. Membranes were blocked for 1 hour in milk block solution and then incubated overnight at 4°C with primary antibodies for either β-actin (Sigma A5441, 1:2000), IL-1β (R&D AF401NA, 1:800) or caspase-1 (Adipogen AG-20B-0042, 1:1000) diluted in milk block solution. Membranes were then washed 5 times for 5 minutes each in wash buffer (PBS supplemented with 0.5% Tween-20) and incubated with goat anti-mouse HRP-conjugated IgG (GE NA931) diluted 1:5000 (β-actin and Caspase-1) or 1:1000 (IL-1β) in milk block for 2 hours at room temperature. Following additional washes, membranes were incubated in HyGlo spray chemilunescent substrates (Denville Scientific, E2400) for 5 minutes and imaged on a Biorad ChemiDoc MP imaging system.

### Cytokine Analysis

The Cytokine Bead Array Mouse Inflammation kit (BD Biosciences 552364) was used according to manufacturer’s instructions for simultaneous measurement of IL-6, IL-10, CCL2, IFNγ, TNFα, and IL-12p70 in supernatants from stimulated BMDMs. General statistical analysis was performed using GraphPad Prism 4.0 (GraphPad Software, San Diego, CA), using an unpaired Student *t* test. For each experiment, both the standard deviation and the standard error of the mean were calculated. P-values of <0.05 were considered significant.

### Identification of Differentially Expressed Genes by RNA-Seq

Total RNA was extracted from three biological replicates of WT and MyD88−/− BMDMs, either unstimulated or stimulated with Bb at MOI 10:1 or MOI 100:1 for 6 hours. Following stimulation, RNA was isolated using the total RNA isolation kit (Macherey-Nagel) and was used as input for the Ovation RNA-seq V1 kit (NuGen, San Carlos, CA). cDNA output was analyzed for correct size distribution with an Experion Standard Sensitivity RNA chip and quantified using a Qubit Fluorometer (Invitrogen, Carlsbad, CA). Sequencing libraries were produced using the NuGen Encore NGS Library I kit. Libraries were multiplexed and sequenced at The Jackson Laboratory for Genomic Medicine Sequencing Core with an Illumina HiSeq 2500 as 2X50bp pair end reads. RNA-Seq reads from each individual library were mapped with Tophat2 RNA-Seq spliced reads mapper (version 2.0.5) (85) to mouse genome build mm9 with parameter settings adjusted to suit strand-specific pair-end RNA-Seq reads. The mapping result bam files were used as input to the HTSeq high-throughput sequencing data analysis package (86) to quantify the read counts mapped to all genes in UCSC mm9 mouse gene annotation set. The expression levels of genes represented as mapped read counts were normalized using the DESeq2 RNA-Seq analysis package (function: *estimateSizeFactor*) (87). Genes were considered expressed if the number of reads was above the 25^th^ percentile for the normalized data set. For quality control, only replicates with Pearson correlation coefficient above 0.9 on their FPKM values were considered (**Figure S5**). Expressed genes were then further analyzed for differential gene expression using the DEseq2 package with FDR cutoff: 0.1. Differential gene expression was calculated in WT BMDMs stimulated 10:1 with Bb relative to unstimulated WT BMDMs and MyD88−/− BMDMs stimulated 100:1 with Bb relative to unstimulated MyD88−/− BMDMs. Differentially expressed genes (DEGs) were classified as either upregulated or down regulated based on the log2 of the fold change compared to the unstimulated control, which was calculated in R statistical software using package “DESeq2”. Determined DEGs were then separated into five groups based on their expression profiles; WT (all DEGs in WT BMDMs), MyD88−/− (all DEGs in MyD88−/− BMDMs), MyD88-dependent (all DEGs in WT but not MyD88−/− BMDMs), MyD88-independent (all DEGs in both WT and MyD88−/− BMDMs), and MyD88-exclusive (all DEGs in MyD88−/− but not in WT BMDMs).

### Identification of Enriched Transcription Binding Sites and Master Regulator Analysis

Transcription factor binding sites in promoters of differentially-expressed genes were analyzed using known DNA-binding motifs described in the TRANSFAC library (88), release 2017.2, available in the GeneXplain software (http://genexplain.com). Binding site enrichment analysis for each one of our sets of DEGs was carried out as part of a GeneXplain dedicated workflow. The background consisted of 300 mouse house-keeping genes and the TRANSFAC mouse Positional Weight Matrices PWM (motifs) for binding site prediction with p-value<0.001 score cutoff. Promoters were extracted by the workflow with a length of 600 bp (−500 to +100) and an enrichment fold of 1.0.

Master regulatory molecules were searched for in signal transduction pathways upstream of the identified transcription factors. The GeneXplain workflow available for this analysis was used in conjunction with the GeneWays database. Parameters set included a maximum radius of 10 steps upstream of the transcription factor nodes, the DEG lists from the respective group as context genes and a z-score cutoff of 1.0. All transcription factors and master regulators used in the network analysis had confirmed expression in respective conditions using the total gene expression lists from the RNA-sequencing data set.

### Gene Ontology (GO) Enrichment Analysis

A Gene Ontology (GO) enrichment analysis was performed for the different sets of DEGs, transcription factors, and master regulators using the TRANSPATH (89) database through GeneXplain software. Input sets were the DEGs, transcription factors, or master regulators from either the MyD88-dependent, MyD88-independent, or MyD88-privative groups. Focus was directed to the Gene Ontology (GO) biological processes output. GO biological processes related to Bb uptake, inflammation, and chemotaxis were identified by first reviewing previous studies for any genes involved in response to Bb relating to these phenotypes. Enrichment analysis was performed on these genes to identify GO biological processes that hit at least 60% of the genes on the list, generating a list of relevant GO biological processes. Then an intersection was performed between the list of GO biological processes identified using our DEG, transcription factor, or master regulator lists, and the GO biological processes identified from the relevant genes. Heat maps of expressed genes hits in each biological process were done in R statistical software using package “ggplots”.

### Network Reconstruction and Network Analysis

Networks were constructed joining the three identified layers on the networks: DEGs, transcription factors, and master regulators. The subnetworks were extracted from identified master regulators of interest. From the MyD88-dependent master regulator group, effort was directed on linking MyD88 with transcription factors that had binding sites in the promoter regions of the MyD88-dependent DEGs enriched in uptake biological processes. These transcription factors were identified using the TRANSPATH database with the enriched DEGs of interest as input. The output list of transcription factors was intersected with the list of transcription factors that were only expressed in WT BMDMs. Networks were assembled and analyzed using Cytoscape software (90). To extract the desired subnetworks, we used OCSANA (91) within the BiNOM plugin (92) in Cytoscape 2.8.3. MyD88 was considered as a source node and transcription factors from the intersected list as target nodes. For MyD88-privative chemotaxis subnetwork construction the same analysis pipeline was applied. MyD88-privative master regulators significantly enriched in chemotaxis were used as source nodes and MyD88-privative transcription factors enriched in chemotaxis were used as targets.

## ACKNOWLEDGEMENTS

The authors would like to thank Anna Allard, Morgan Ledoyt and Dr. Ashley Groshong for their contribution and support to execution of the experiments. The authors would also like to thank Dr. Reinhard Laubenbacher for resources provided when developing computational data. This work was funded by both the National Institutes of Health and Connecticut Children’s Medical Center.

## AUTHOR CONTRIBUTIONS

Sarah Benjamin (Figures 1-7): Conceptualization, Formal Analysis, Investigation, Methodology, Validation, Visualization, Original Draft Preparation, Review and Editing Kelly Hawley (Figures 1-3, 7): Methodology, Project Administration, Supervision, Review and Editing

Paola-VeraLicona(Figures4-6):Conceptualization, Formal Analysis, Project Administration, Supervision, Validation, Review and Editing

Carson J. LaVake (Figures 1-7): Investigation, Methodology, Review and Editing

Jorge L. Cervantes (Figures 4-6): Investigation

Rachel Burns: (Figure 1): Formal Analysis, Review and Editing

Oscar Luo: (Figures 4-6): Formal Analysis, Software

Yijun Ruan: (Figures 4-6): Resources

Melissa J. Caimano (Figures 1-3, 7): Methodology, Project Administration, Validation, Review and Editing

Justin D. Radolf (Figures 1-7): Conceptualization, Project Administration, Review and Editing

Juan C. Salazar (Figures 1-7): Conceptualization, Funding Acquisition, Project Administration, Resources, Supervision, Original Draft Preparation, Review and Editing

## DECLARATION OF INTERESTS

The authors have no interests to declare.

## SUPPLEMENTAL FIGURE CAPTIONS

**Figure S1:**
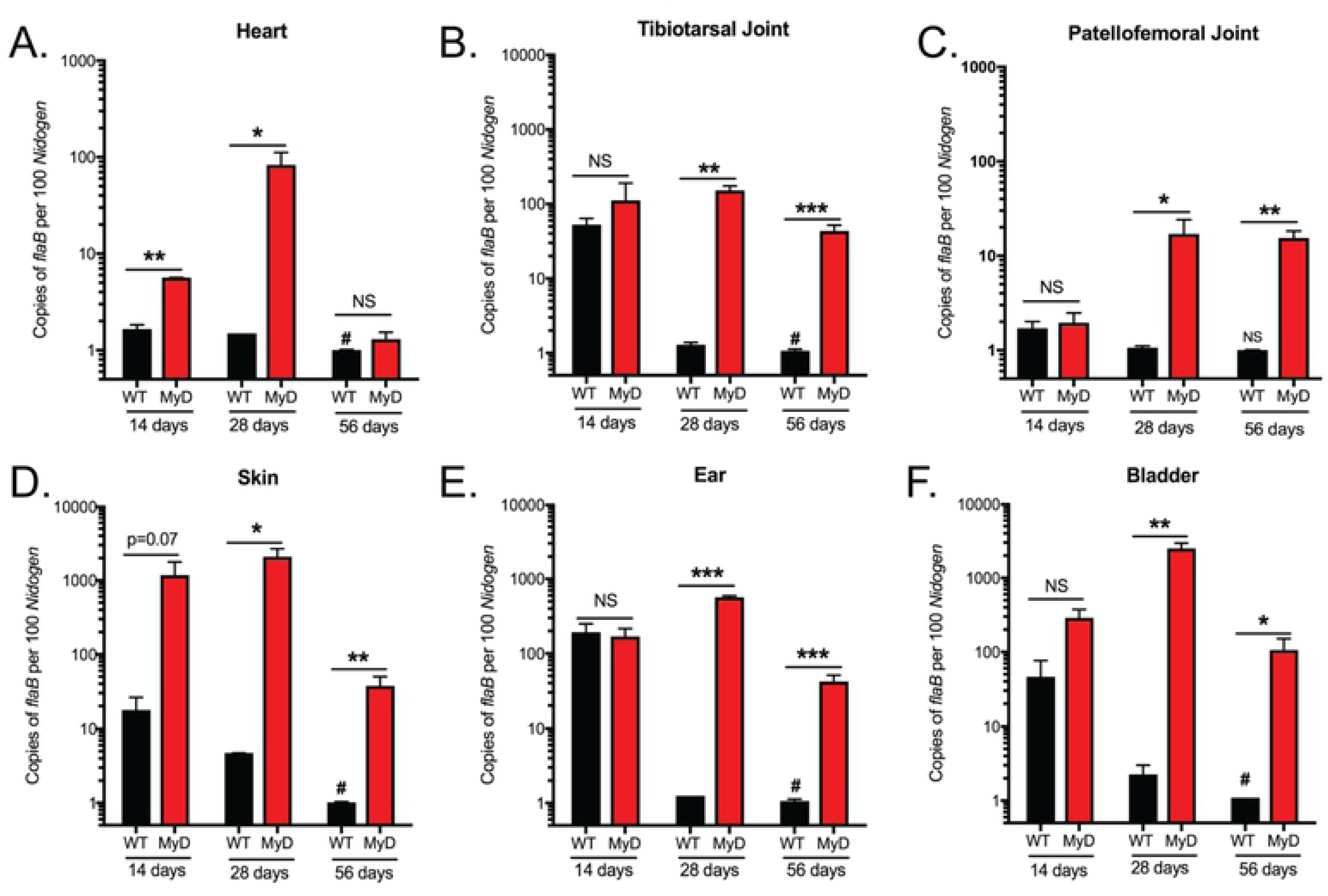
MyD88−/− mice show increased Bb burdens 28 and 56 DPI. (A-F) Bb burdens in selected mouse tissues as determined by RT-PCR. Mice were syringe-inoculated with 10^5^ Bb914 and sacrificed 14, 28 or 56 DPI. DNA was isolated from WT (black bars) and MyD88−/− (red bars) mice. Copies of *flaB* were normalized to *Nidogen* amplified from total tissue DNA in heart (A), tibiotarsal joint (B), patellofemoral joint (C), skin (D), ear (E), and bladder (F). n=3-5 mice per group *p-value<0.05, **p-value<0.01, NS=not significant, ^#^p-value<0.05 using Mann-Whitney test against WT 14 DPI burden values

**Figure S2:**
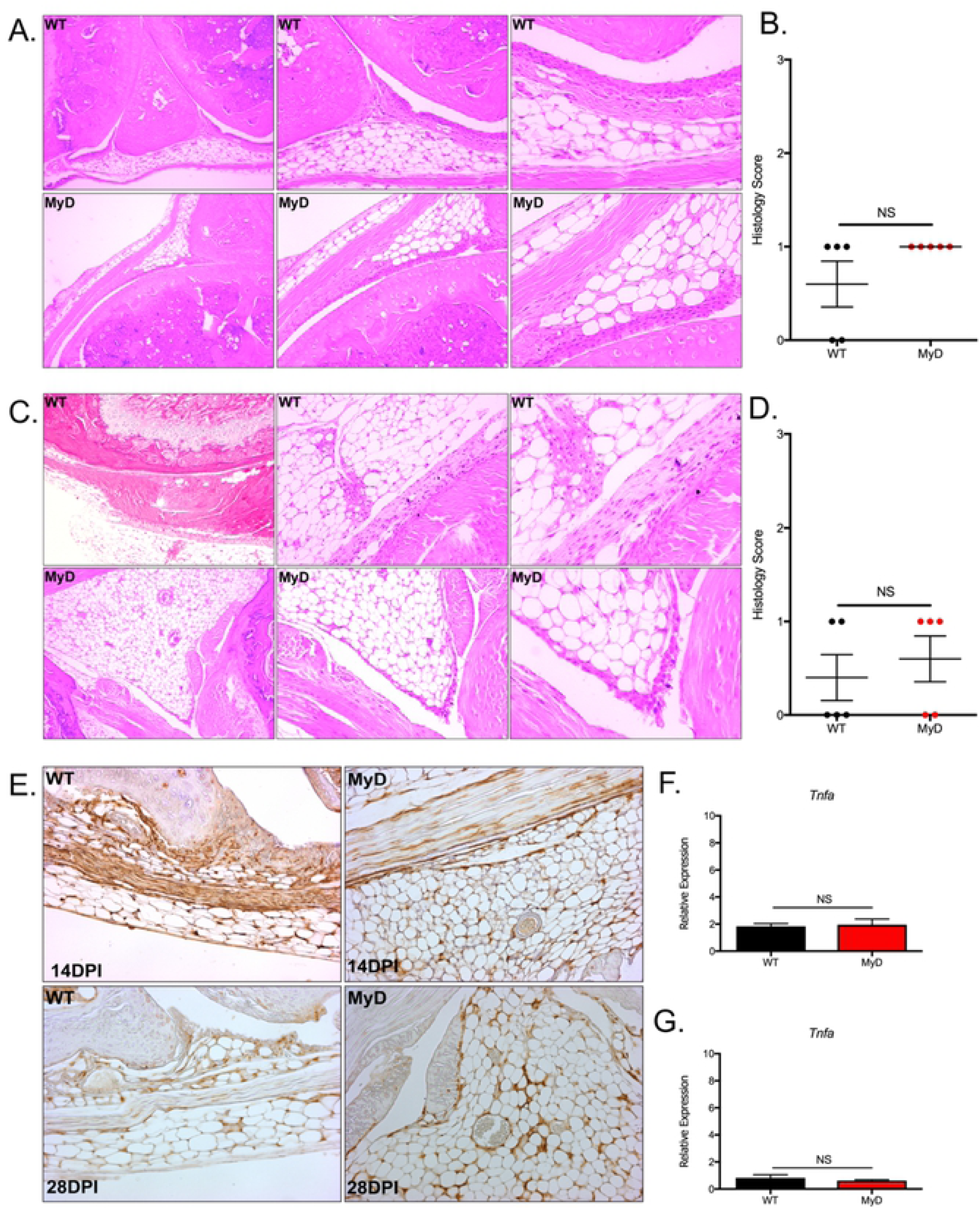
MyD88−/− mice show comparable inflammation and macrophage infiltrate in patellofemoral joint tissue 14- and 28-days post Bb-infection. (A and C) Representative images of H&E staining of patellofemoral tissue sections from WT and MyD88−/− mice syringe-inoculated with 10^5^ Bb at 14 DPI (A) and 28 DPI (C). Images are of increasing magnification from left to right (10x, 20x, 40x). Sections are 5μm. (B and D) Compilation of inflammation scores for WT (black dots) and MyD88−/− (red dots) mice at 14 DPI (B) and 28 DPI (D) N=5 mice per group. (E) Iba-1 immunohistochemistry of patellofemoral joint tissue sections from Bb-infected WT and MyD88−/− mice, magnification = 20x. Sections are 5μm. (F and G) Compilation of qRT-PCR *Tnfa* gene amplifications from patellofemoral joint tissue RNA. Total RNA was isolated from Bb-infected WT and MyD88−/− mice 14 (F) and 28 (D) days post infection. Gene amplification values were normalized to *Gapdh*. *p-value<0.05, **p-value<0.01, NS=not significant.

**Figure S3:**
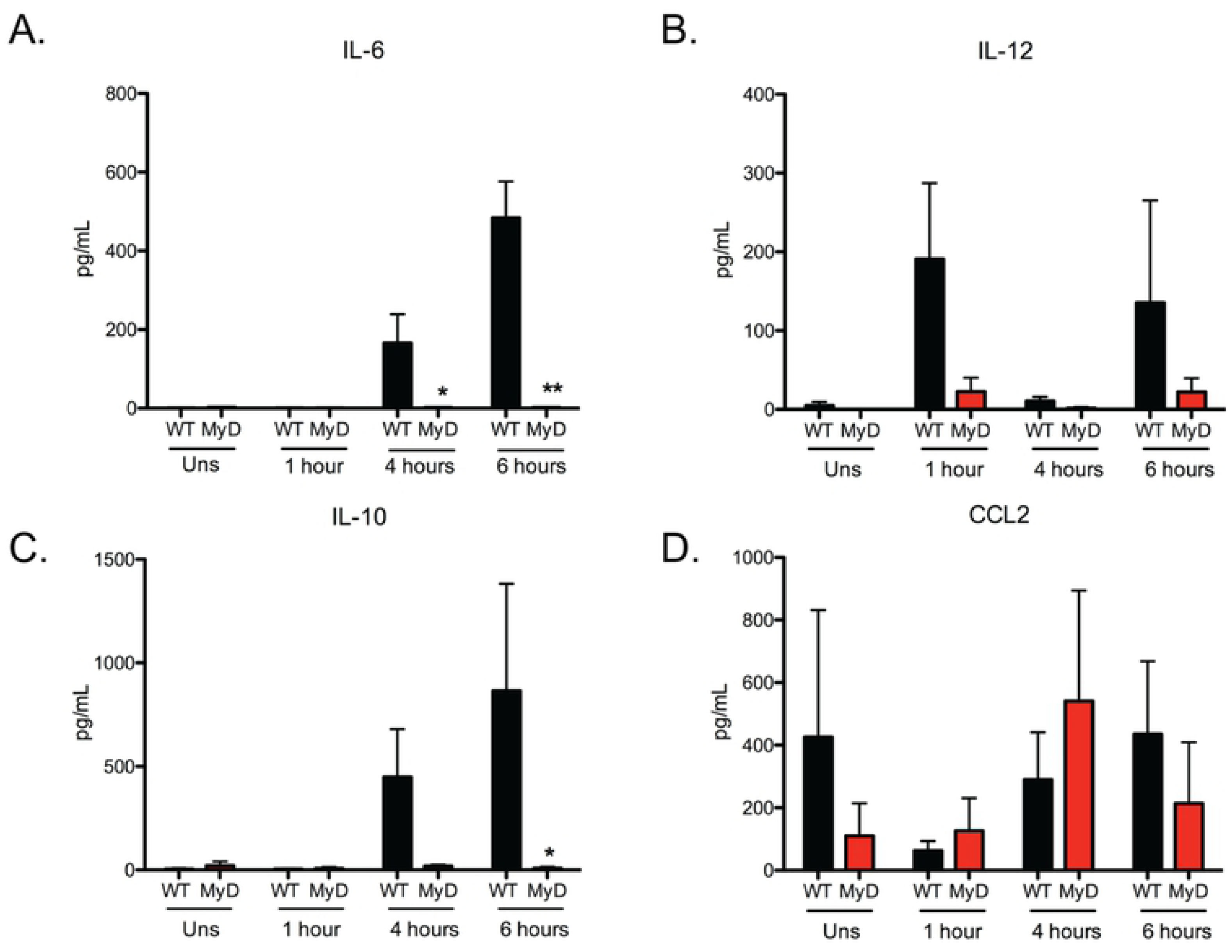
MyD88−/− BMDMs stimulated with Bb show abrogated cytokine production. (A-D) Quantification of IL-6 (A), IL-12 (B), IL-10 (C) and CCL2 (D) proteins in supernatant from WT and MyD88−/− (MyD) BMDMs stimulated with Bb at MOI 10:1 for 1, 4 or 6 hours. N= 3 mouse BMDM per genotype *p-value<0.05, **p-value<0.01, ***p-value<0.001, NS=not significant

**Figure S4:**
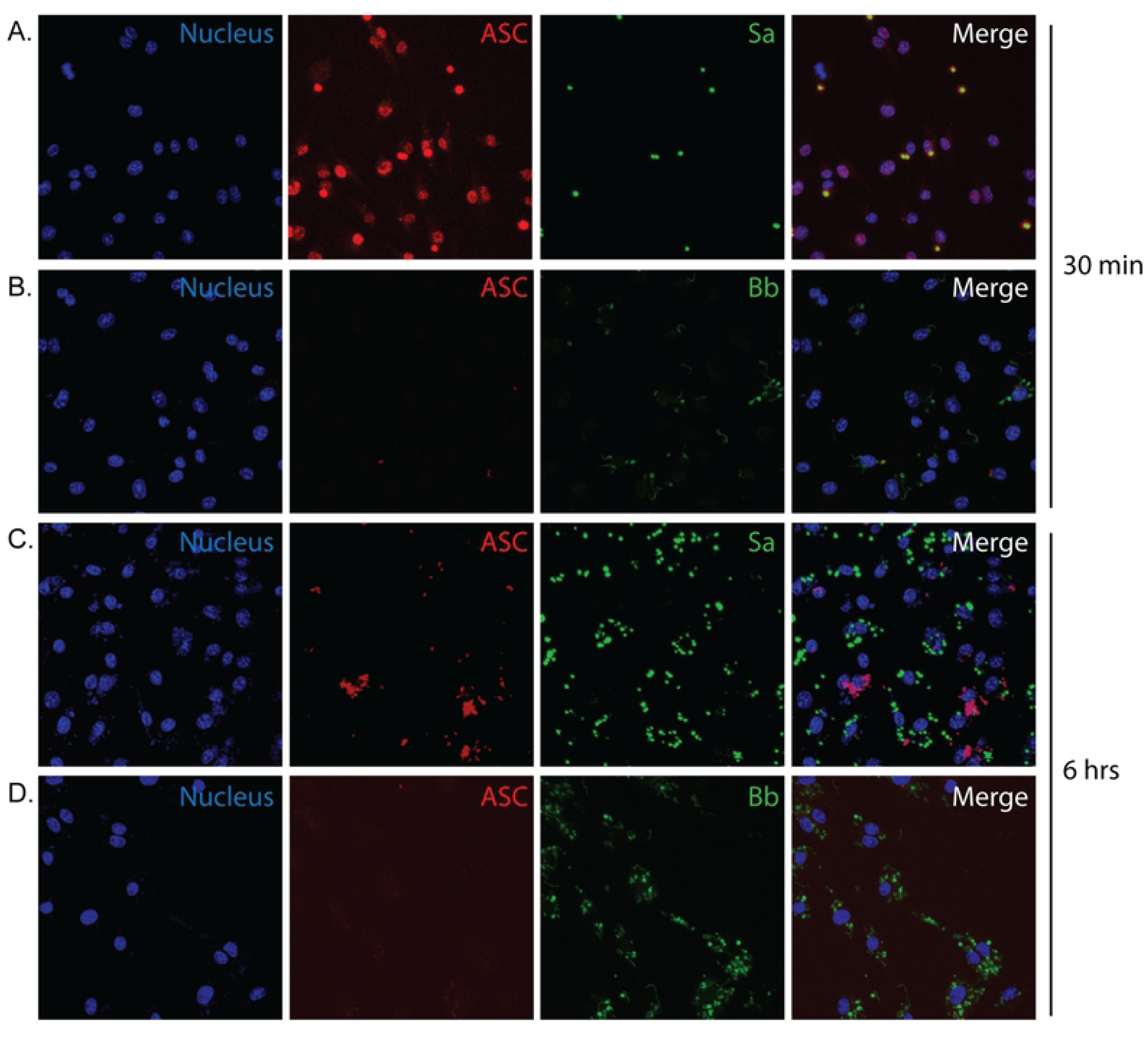
Bb does not induce ASC formation in BMDMs. (A-D) Confocal images (40x) of BMDMs stimulated with either Bb (B and D) or Sa (A and C) for 1 hour (A-B) or 6 hours (C-D). Blue is nucleus, red is ASC and green is the bacteria species.

**Figure S5:**
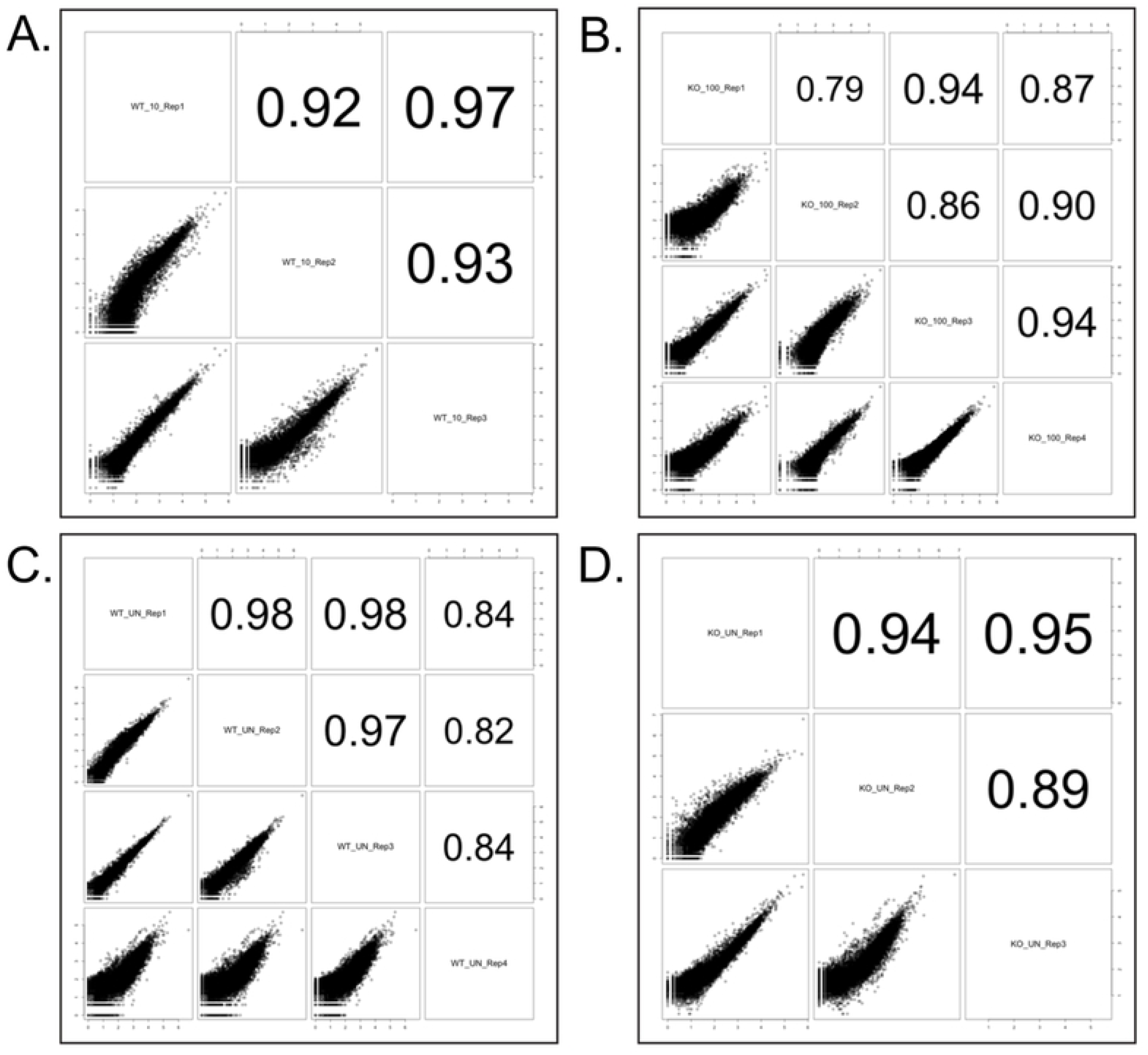
Correlation of reads of RNA sequencing data. (A-D) Correlation of reads of RNA isolated from Bb-stimulated BMDMs. WT BMDMs were stimulated at an MOI 10:1 for 6 hours (A). MyD88−/− BMDMs were stimulated at an MOI 100:1 for 6 hours (B). To calculate differential expression, reads from the stimulated samples were normalized to unstimulated BMDMs of the same genotype; WT (C) or MyD88−/− (D).

**Figure S6:**
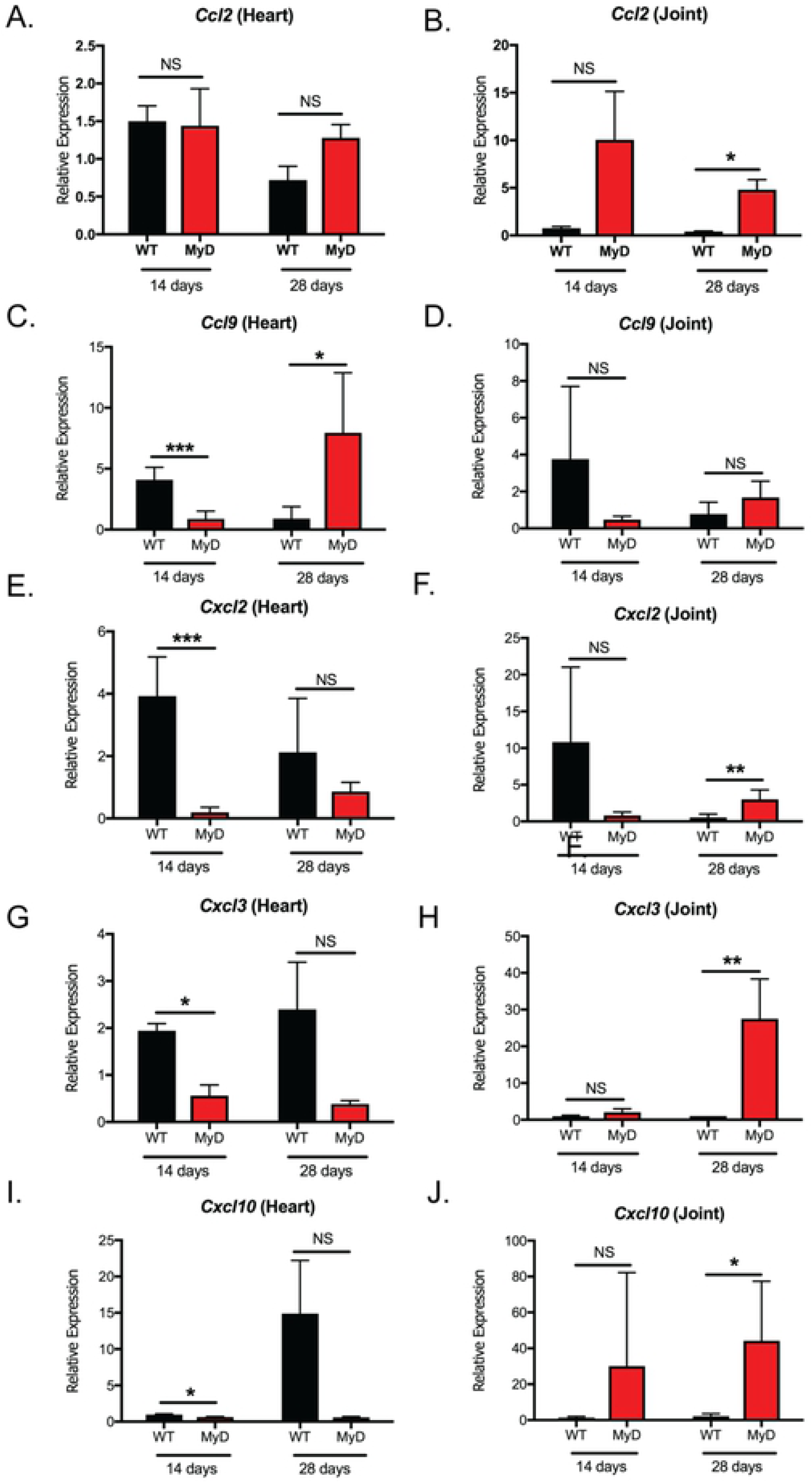
MyD88-independent chemokines are produced in MyD88−/− mice during *in vivo* infection. (A-J) Transcript analysis of Bb-infected WT (black bars) and MyD88−/− (red bars) heart or patellofemoral joint tissue RNA. Total RNA was isolated 14 and 28 DPI. Transcript production of *Ccl2* (A-B), Ccl9 (C-D), *Cxcl2* (E-F), *Cxcl3* (G-H) and *Cxcl10* (I-J) were measured by qRT-PCR from total tissue RNA. Gene amplification values were normalized to *Gapdh*. *p-value<0.05, **p-value<0.01, ***p-value<0.001, NS=not significant

## SUPPLEMENTAL FILES

**Supplemental File 1:** Gene ontology analysis of differentially expressed genes.

**Supplemental File 2:** Gene ontology analysis of identified transcription factors.

**Supplemental File 3:** Gene ontology analysis of identified master regulators.

**Supplemental File 4:** Identified differentially expressed genes.

**Supplemental File 5:** Identified transcription factors.

**Supplemental File 6:** Identified master regulators.

